# From Initial Formation to Developmental Refinement: GABAergic Inputs Shape Neuronal Subnetworks in the Primary Somatosensory Cortex

**DOI:** 10.1101/2022.10.23.513371

**Authors:** Jui-Yen Huang, Michael Hess, Abhinav Bajpai, Xuan Li, Liam N Hobson, Ashley J Xu, Scott J Barton, Hui-Chen Lu

## Abstract

Neuronal subnetworks, also known as ensembles, are functional units formed by interconnected neurons for information processing and encoding in the adult brain. Our study investigates the establishment of neuronal subnetworks in the mouse primary somatosensory (S1) cortex from postnatal days (P)11 to P21 using *in vivo* two-photon calcium imaging. We found that at P11, neuronal activity was highly synchronized but became sparser by P21. Clustering analyses revealed that while the number of subnetworks remained constant, their activity patterns became more distinct, with increased coherence, independent of cortical layer or sex. Furthermore, the coherence of neuronal activity within individual subnetworks significantly increased when synchrony frequencies were reduced by augmenting GABAergic activity at P15/16, a period when the neuronal subnetworks were still maturing. Together, these findings indicate the early formation of subnetworks and underscore the pivotal roles of GABAergic inputs in modulating S1 neuronal subnetworks.

## Introduction

For the brain to perform complex information processing, its circuits organize into small functional units called subnetworks or ensembles. Neurons in the same subnetwork exhibit recurrent coordinated activity and are consistently recruited by specific sensory stimuli, motor programs, or cognitive states ^1–9^. Several studies have shown that neurons from different subnetworks, encoding different types of information, are spatially intermingled and distributed across large brain volumes ^1,2,4,7^. Despite their wide topographical distribution, neurons within each subnetwork also co-activate with activity patterns that resemble externally evoked activity patterns when external inputs are weak or absent ^8,10–12^. These findings suggest that a subnetwork is a pre-configured functional unit ready to receive or integrate information input. Currently, it is unclear when such functional subnetworks assemble, and what factors refine their functional connectivity during brain development.

Two-photon calcium imaging has been used to visualize neuronal activity at population levels in the developing brain ^13–22^. These studies typically evaluated activity patterns with two metrics: synchrony frequency, which measures the simultaneous burst activation of neurons, and the Pearson correlation coefficient, which quantifies the degree of correlated activity between neuron pairs ^15,17–20^. During the first postnatal week, both glutamatergic and GABAergic neurons in the developing rodent primary somatosensory cortex (S1) exhibit high levels of synchronization and correlation ^15–22^. Three distinct phases of activity patterns have been identified in S1 layer (L) 4 neurons during postnatal day (P)1 to P11: patchwork-type (P1-P7), synchronized (P7-P11), and sparse (after P11) ^15,17^. From P11 to P21, L2/3 neurons exhibit highly synchronous and correlated activation patterns before transitioning to sparser patterns^18,19,21,23^.

During this postnatal period, particularly after P7, both excitatory and inhibitory circuits in S1 cortex become more established and acquire greater strength ^24–27^. Around P9, recurrent excitatory connectivity in L4 glutamatergic neurons increases ^26,27^, while the number of inhibitory synapses on glutamatergic neurons gradually rises from P9 to P28 ^24,25^. Between P10 and P20, the frequency of spontaneous inhibitory postsynaptic currents (sIPSCs) in L2/3 excitatory neurons increases, while the frequency of spontaneous excitatory postsynaptic currents (sEPSCs) decreases ^25^, suggesting that inhibition increases more than excitation. Given that the maturation of gamma-aminobutyric acid (GABA)ergic neurons aligns with the desynchronization of population activity, we hypothesized that increasing inhibitory inputs contribute to the transition from synchronized activity to sparse coding, thereby influencing subnetwork assembly in the S1 cortex.

In this study, we aimed to investigate the timing and mechanisms of S1 cortical neuron subnetwork assembly and explore the role of GABAergic inputs on subnetwork refinement.

Using two-photon calcium imaging, we monitored the activity of S1 upper cortical layer (L2-4) neurons in awake neonatal mice from P11-P21. To identify neuronal subnetworks, we applied several clustering algorithms, including k-means clustering, a community detection method, and density-based spatial clustering of applications with noise (DBSCAN)^11,16,28^, to identify neurons exhibiting similar activity patterns and to group them into individual subnetworks. All these clustering analyses indicated the presence of neuronal subnetworks from P11, the earliest postnatal age we imaged. Notably, the cohesiveness of activity within these subnetworks increased significantly from P11 to P21, regardless of cortical layer or sex. These findings highlight the early assembly of S1 neuronal subnetworks and the subsequent refinement in enhancing subnetwork coherence. Artificially enhancing GABAergic neuronal activity at P15/P16 reduced synchrony frequency and increased subnetwork coherence. This latter result provides strong support for the contribution of GABAergic transmission in subnetwork maturation.

## Results

### Spontaneous neuronal activity decreased and desynchronized in both L2/3 and L4, irrespective of sex, from P11 to P21

To elucidate when S1 neuronal subnetworks assemble during postnatal brain development, we conducted *in vivo* two-photon calcium imaging with the red calcium sensor jRGECO1a to monitor neuronal activity in awake-behaving mouse pups. We injected an AAV carrying jRGECO1a into S1 at P1-P2 via stereotaxic intracranial injections and installed a cranial window at P8-P9 (Fig. 1A, Fig. S1A-B). We imaged pups in a head-restrained configuration (Fig. S1C-D) at P11, P13, P15, P18, and P21. Sequential imaging of a 505 x 505 µm field at different cortical depths was conducted to capture neuronal activity in the upper S1 cortical layers, L2/3 and L4. The cranial window installed on P8-9 cleared within two days and remained so until the final imaging day, P21 (Fig. 1B, Fig. S1F upper panel). Guided by vasculature patterns, we imaged the same cortical areas over the whole imaging period (Fig. S1F lower panel, Videos S1, S2, and S3).

**Fig. 1.**
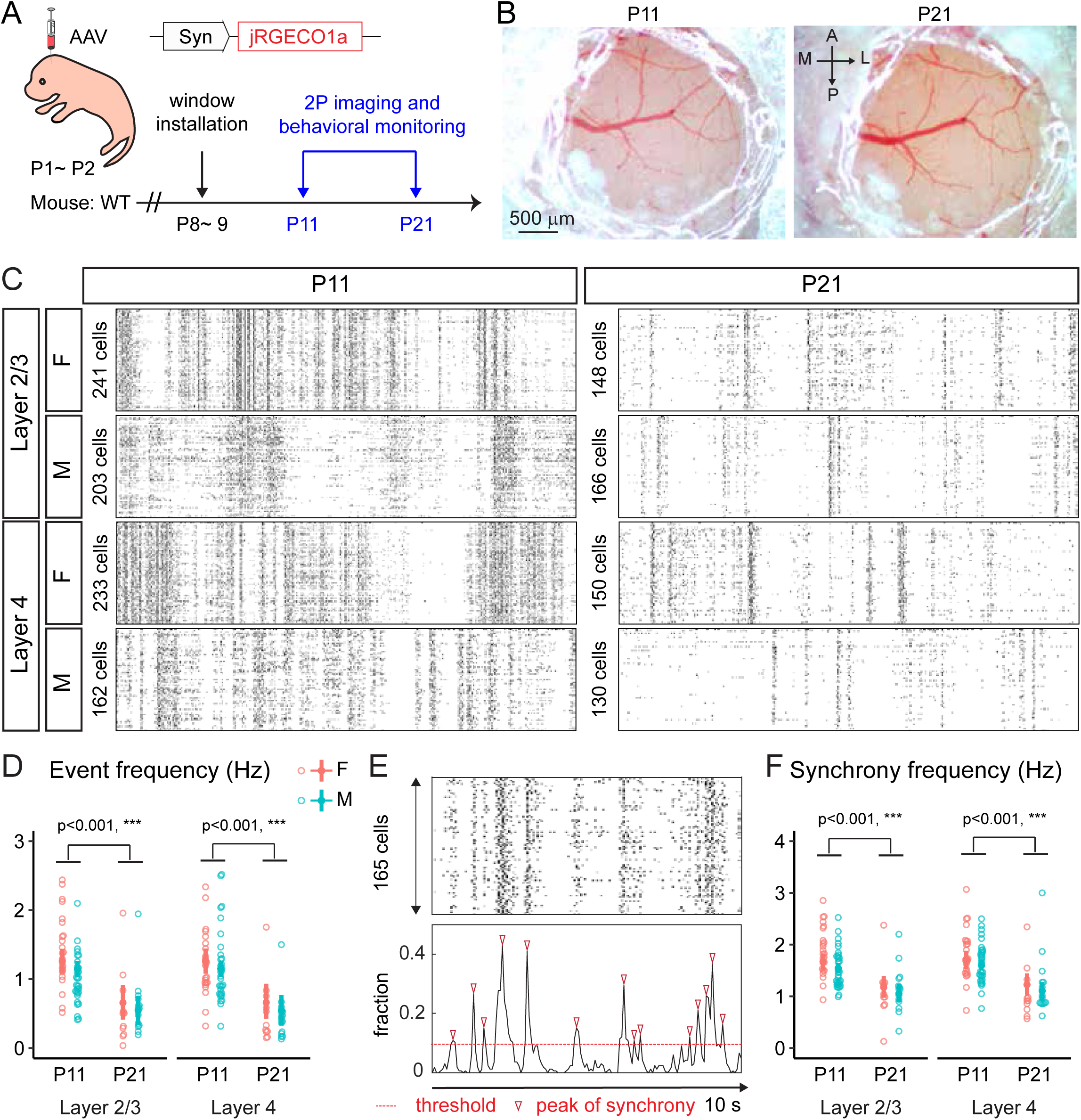
Spontaneous neuronal activity gradually decreased and desynchronized in both L2/3 and L4, irrespective of sex, with increasing age. (A) Schematic of the experimental timeline. Mice received injections of AAV carrying jRGECO1a (AAV9-Syn.NES-jRGECO1a) at P1-2, underwent cranial window installation at P8-9, and were subsequently imaged from P11 to P21. (B) Representative bright field images showing the cranial window implanted at P9, which remained optically transparent throughout the study. A, anterior; P, posterior; M, midline; L, lateral. (C) Example raster plots of spontaneous neuronal activity. (D) Event frequency of L2/3 and L4 neurons. (E) Upper panel: raster plot of L2/3 neurons from a P11 female pup. Lower panel: cumulative fraction of active neurons over time corresponding to the upper panel. Synchronous events were identified when the fraction of recruited neurons exceeded the threshold. (F) Frequency of synchronous events in L2/3 and L4 neurons. Numbers of experimental mice. P11, 14 female (F), 20 male (M); P21, 6F, 11M. Data are presented as mean ± 95% confidence intervals derived from the bootstrapping output.

Raw calcium imaging datasets were preprocessed using Suite2P and FISSA for movement and neuropil correction, followed by the detection of calcium transients to estimate neuronal firing rates (Fig. S2A-D, see in STAR Methods). To reveal cortical activity during the default mode, we employed cameras to simultaneously record body and facial movements of the mouse during calcium imaging. This allowed us to identify periods when the mouse pups were in a quiescence status, with minimal body movement, for subsequent calcium data analysis (Fig. S2E). Given the nested nature of the experimental design ^29–31^, such as imaging the same mice multiple times, we primarily employed linear mixed-effects (LME) models for multilevel statistical analysis ^32^. In cases where the data did not exhibit nesting, we used linear regression (LM) models to assess statistical significance instead (see in STAR Methods).

To investigate sex- or layer-dependent effects on spontaneous neuronal activity during cortical development, we first compared data from P11 and P21 for males and females in both L2/3 and L4. We observed a notable developmental decline in event frequency from P11 to P21 in L2/3 and L4, a consistent trend in both sexes (Fig. 1C, D). To quantify overall network activity, we computed synchrony events (Fig. 1E; see in STAR Methods). The results showed a significant decrease in synchrony frequency by P21, regardless of cortical layer or sex (Fig. 1F). Next, we evaluated the correlated activity between neuronal pairs by calculating the Pearson correlation coefficient of normalized fluorescent traces from all neuronal pairs (Fig. S2F). Our analysis revealed a significant decrease in the mean correlation coefficient from P11 to P21 in females, but not in males (Fig. S2F). Furthermore, L2/3 neurons showed higher correlated activity compared to L4 neurons at P11, regardless of sex (Fig. S2F). Because the Pearson correlation captured only the linear relationships between calcium responses of paired neurons, we also calculated mutual information (MI) to quantify the amount of shared information between neuronal calcium dynamics (Fig. S2G, see in STAR Methods). Corroborating the Pearson correlation findings, MI analysis shows a significant decrease in MI at P21 compared to younger ages in both sexes and cortical layers. Notably, at P21, male neurons displayed significantly higher MI values than female neurons, regardless of cortical layer. Additionally, L4 neurons exhibited lower MI values compared to L2/3 neurons at both P11 and P21, independent of sex.

In summary, our developmental calcium imaging studies revealed that spontaneous neuronal activity decreased and became desynchronized in both L2/3 and L4 cortical neurons from P11 to P21, irrespective of sex. While the overall activity patterns showed a similar trend, our analyses identified subtle but significant sex- and layer-dependent differences in the extent and nature of the correlated activity.

### Subnetworks are formed by P11 and become matured at P21 with increased coherence in activity patterns

We next investigated the development of cortical neuronal subnetworks between P11 and P21. To find neurons exhibiting similar activity patterns, we employed an unsupervised k-means clustering algorithm ^11,16^. This algorithm clustered the similarity matrix (covariance matrix) generated from frame-by-frame analysis of calcium events (Fig. S3A-B). The order of neurons and calcium events were rearranged according to their event clusters and visually represented with color coding (Fig. 2A, B; see in STAR Methods). Subsequently, neurons were assigned to the activity event cluster to which they made the greatest contribution, thereby revealing the neuronal subnetwork (Fig. 2C).

**Fig. 2.**
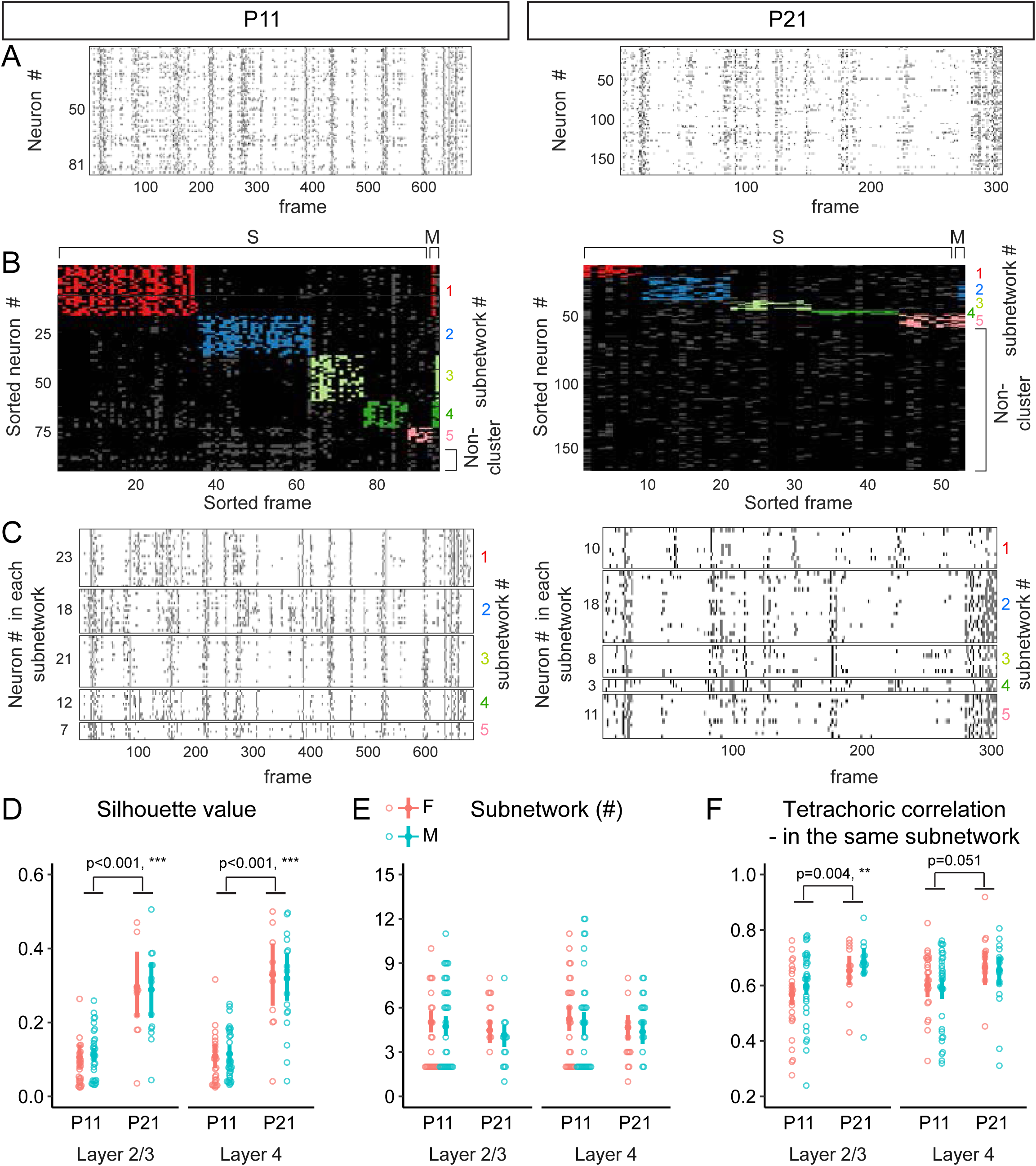
Developmental increase in subnetwork coherence. (A) Raster plots showing spontaneous neuronal activity during quiescence at P11 (left) and P21 (right). (B) Re-sorted raster maps based on neuronal subnetwork (y-axis) and event cluster (x-axis). Five distinct neuronal subnetworks (color-coded) were identified at P11 and P21 after assigning neurons to the event clusters in which they participated the most. Abbreviations: S, events associated with a single event cluster; M, events associated with multiple event clusters. Event clusters without significant neuronal participation were excluded. (C) Raster plots of neurons within each identified subnetwork, with neurons grouped according to their subnetwork identities (color-coded numbers). (D) Silhouette value of event cluster. (E) Number of neuronal subnetworks. (F) Tetrachoric correlation coefficient of neuronal pair within the same subnetwork. Numbers of experimental mice. P11, 14F, 20 M; P21, 6F, 10M. Data are presented as mean ± 95% confidence intervals derived from the bootstrapping output.

To evaluate the quality of event clusters, we inspected the silhouette value (Fig. 2D), which serves as a metric for evaluating the similarity of a point to its own cluster (cohesion) and individuality compared to other clusters (separation). In other words, a higher silhouette value indicates greater similarity to other members within the same cluster and/or lower similarity to members of other clusters. After assigning neurons to the activity clusters to which they contributed most significantly (Fig. 2B-C), we found that neuronal subnetworks were already present at P11 and that a similar number of subnetworks was found at P21 (Fig. 2E). Interestingly, our data analysis revealed a substantial increase in silhouette value from P11 to P21, spanning both cortical layers and sexes (Fig. 2D).

To validate the efficacy of the k-means clustering algorithm in revealing subnetworks, we computed the tetrachoric correlation coefficient (tCC)—a suitable measure for binary data analogous to the Pearson correlation coefficient ^33^— for all neuronal pairs, regardless of their subnetwork identities. This analysis involved calculating the tCC for both neuronal pairs within their specific subnetworks (inside) and pairwise for neurons between subnetworks (outside). Notably, the tCC of neuronal pairs within the same subnetworks at P11 and P21 was consistently higher than that of pairwise comparisons between neurons from different subnetworks (Fig. S3C). These results demonstrate the robustness of the clustering algorithm in grouping neurons based on the similarity of their activity patterns. Additionally, we observed that L2/3 neurons had a significantly heightened tCC at P21 compared to P11, regardless of sex (Fig. 2F), while a similar trend was observed in L4 neurons, without reaching statistically significance (p = 0.051).

To evaluate whether the number of neurons and time intervals used in the analysis could influence the identification of subnetworks, we examined their respective correlations with the number of identified subnetworks. The Spearman correlation coefficient for the number of neurons included in the clustering analysis was ρ = -0.063 (Fig. S3D) and for the time intervals used, ρ = 0.233 (Fig. S3E). These results indicate that both the number of neurons and time intervals had minimal impact on subnetwork identification.

### Changes in subnetwork configuration and topographical distribution: a comparison between P11 and P21, from high participation to low and from segregation to intermingling

We next explored whether the higher correlated activity among neurons within the same subnetwork is maintained by removing neurons that exhibit lower correlated activity. We classified neurons based on their involvement in event clusters. For those participating in clusters, we examined whether they exclusively engaged in a single event cluster or were part of multiple event clusters. With these characteristics, neurons were separated into the following three categories: single-cluster neurons, non-cluster neurons, and multiple-cluster neurons. Our findings revealed a decrease in the percentage of single-cluster neurons with age (Fig. 3A), accompanied by a rise in the percentage of non-cluster neurons (Fig. 3B). The percentages of multiple-cluster neurons remained similar between P11 and P21 (Fig. 3C). This pattern of configuration was consistent across both L2/3 and L4 layers and between sexes (Fig. 3A-C).

**Fig. 3.**
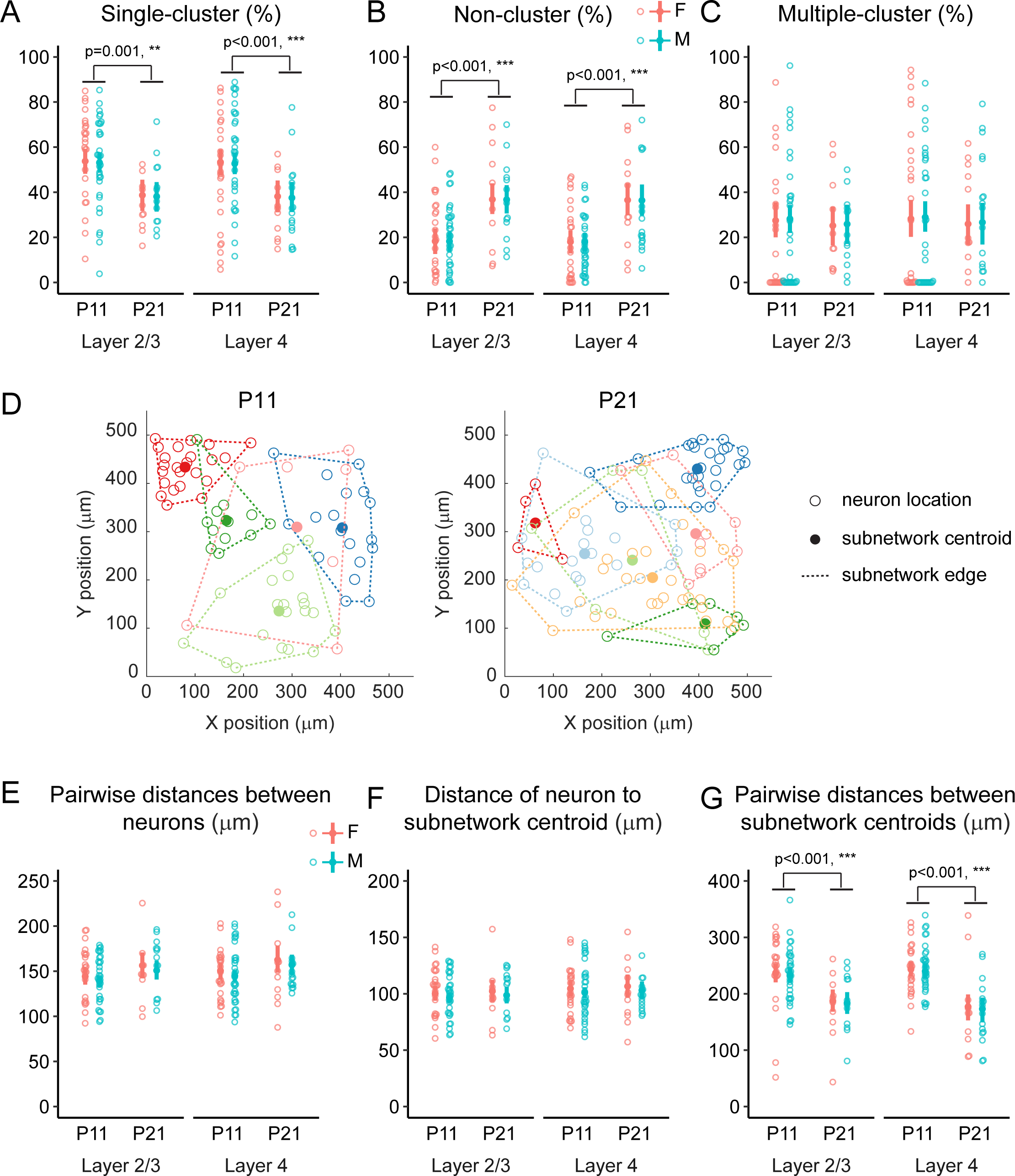
Subnetwork configuration and topographical distribution. (A) Percentage of neurons participating in single event clusters. (B) Percentage of neurons not participating in any event cluster. (C) Percentage of neurons participating in multiple event clusters. (D) Neuronal topographical map showing the distribution of neurons within each subnetwork (color-coded). Open circles represent the locations of individual neurons, while filled circles represent the centroids of the corresponding subnetworks. The subnetwork edges, estimated using the convex hull method, are indicated by dashed lines. Neurons not grouped into subnetworks are not plotted. (E) Pairwise distances among neurons within the same subnetwork. (F) Distances of neurons to their respective subnetwork centroids. (G) Pairwise distances between subnetwork centroids. Animal numbers are the same as in Fig. 2. Data are presented as mean ± 95% confidence intervals derived from the bootstrapping output.

Previous studies have demonstrated that neurons within the same subnetwork are spatially intermingled and distributed across large brain volumes in adult mice ^1,2,4^. Here we investigated the topographical distribution of each neuron based on their subnetwork identification and determined if their distribution patterns changed with age (Fig. 3D). To determine whether the physical size of neuronal subnetworks changes with age, we calculated the pairwise distance between neurons within the same cluster and the distance of each neuron to its cluster centroid. We found no significant differences between P11 and P21 in either measure within the same cortical layers (Fig. 3E, F). To evaluate the relative distributions among subnetworks, the pairwise distances between subnetwork centroids were quantitatively compared. We observed a significant reduction in these distances from P11 to P21 for both cortical layer and sexes (Fig. 3G).

In summary, our results demonstrate that neuronal subnetworks in the S1 upper cortical layer are assembled as early as P11, and the number of subnetworks remains consistent between P11 and P21. Notably, by P21, these subnetworks exhibit greater coherence in activity patterns while having fewer neurons within each cluster. Additionally, the spatial distribution of these subnetworks evolves from initially segregated to a more intermingled pattern.

### The refinement of S1 neuronal subnetworks occurs during the 2^nd^ to 3^rd^ postnatal weeks

To examine the developmental trajectory of cortical subnetwork assembly, we analyzed imaging data acquired from age groups P11, P13, P15, P18, and P21, merging data from females and males when no sex-dependent effect was observed. We quantified mean differences among age groups to reveal the developmental transition periods for individual network properties. A developmental decline in spontaneous neuronal event frequency was observed (Fig 4A), with the greatest mean difference occurring between P11 and P18 (Fig. 4A) while synchrony frequency showed a gradual decline from P11 to P21 (Fig. 4B). Pearson correlation coefficients revealed that, in females, L2/3 and L4 neurons showed a gradual decrease in their correlation coefficients from P11 to P21 and P11 to P18, respectively (Fig. S4A). In males, correlation coefficients decreased from P11 to P15, followed by a transient increase at P18. MI analysis revealed a consistent decline in MI values over time in females (P11-P21) and males (P11-P15), with females exhibiting higher MI values than males at P21(Fig. S4B). Furthermore, L2/3 neurons consistently exhibited higher MI values than L4 neurons across all age groups, regardless of sex or cortical layer (Fig. S4B). Thus, both pairwise Pearson correlation and MI analysis identified similar developmental trends, with subtle but significant cortical layer- and sex-specific changes.

**Fig. 4.**
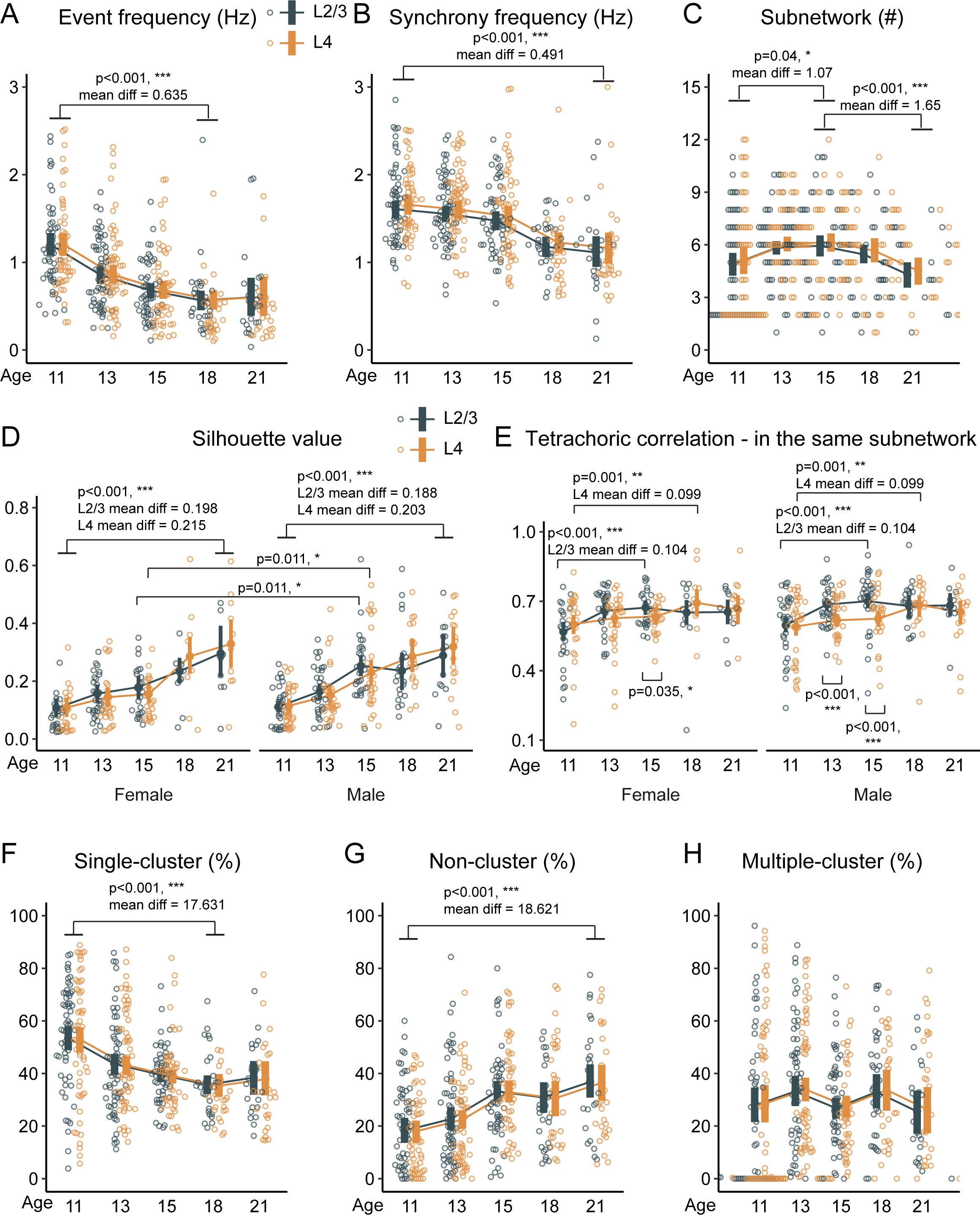
Developmental trajectory of cortical subnetwork assembly during early postnatal stages. This figure summarizes the progression of various parameters associated with cortical subnetwork development from P11 to P21. (A) Event frequency. (B) Synchrony frequency. (C) Subnetwork number. (D) Silhouette value of event clusters. (E) Tetrachoric correlation of neurons within the same subnetwork. (F) Percentage of neurons participating in a single event cluster. (G) Percentage of neurons not involved in any event cluster. (H) Percentage of neurons participating in multiple clusters. Data from both sexes were combined in panels where no sex-dependent effects were observed to streamline the presentation. The age groups with the largest mean differences are annotated to emphasize key developmental transitions. The same animals were used for all figures, and datasets not identified by the k-means clustering algorithm as part of a subnetwork were excluded from the subnetwork-based analysis. Numbers of experimental mice: P11, 14 female (F), 20 male (M); P13, 13F, 18M; P15, 11F, 17M; P18, 5F, 11M; P21, 6F, 11M. Data are presented as mean ± 95% confidence intervals derived from the bootstrapping output.

With the subnetwork analysis, we found a transient but modest increase in the subnetwork number at P15 (Fig. 4C). For event cluster coherences, the silhouette value gradually increased from P11 to P21, regardless of sex and cortical layer (Fig. 4D). At P15, males transiently exhibited significantly higher silhouette values than females (Fig. 4D). Interestingly, as indicated by the tCC of neurons within the same subnetworks, L2/3 neurons reached their maximum subnetwork-based correlation earlier than L4 neurons: the maximum mean difference for L2/3 occurred at P15, while for L4 occurred at P18, regardless of sex (Fig. 4E). The developmental trajectory of subnetwork configuration showed that the percentage of single-cluster neurons gradually decreased from P11 to P18 (Fig. 4F), while the percentage of non-cluster neurons increased from P11 to P21 (Fig. 4G). The percentage of neurons participating in multiple clustering remained consistent throughout the observation period (Fig. 4H).

Regarding the spatial distribution of subnetwork neurons, our analysis suggests that the spatial distribution of subnetworks became more intermingled after P18. We found that the maximum mean difference in physical size indices—specifically, the pairwise distances between neurons and the distance of neurons to their subnetwork centroid—occurred between P13 and P21, regardless of cortical layer or sex (Fig. S4B, C). Notably, the pairwise distances between subnetwork centroids gradually decreased from P11 to P21, with the largest reduction observed in L2/3 and L4 neurons during P11-P21 and P11-P18, respectively (Fig. S4D).

In summary, our results demonstrate that nascent subnetworks undergo a dynamic developmental transformation between P11 to P21. The coherences of neuronal activity patterns, as indicated by increased silhouette values, is significantly enhanced during this period. Interestingly, the number of neurons participating in a single subnetwork declines with age. These data suggest that the 2^nd^ to 3^rd^ postnatal weeks are a critical period for subnetwork development, with P15 serving as a particularly notable transition point. During this developmental transition, cortical neurons intricately enhance their functional connections and form highly correlated subnetworks that become increasingly independent of other subnetworks.

### Acute GABAergic transmission augmentation increases the coherence of cortical subnetworks

Inhibitory inputs from GABAergic neurons in S1 cortex increase substantially from the 2^nd^ to 3^rd^ postnatal week ^24,25^, matching the period we observed the developmental transitions for various subnetwork properties. To investigate whether GABAergic inputs play a role in refining S1 cortical subnetwork properties, we employed a designer receptor exclusively activated by designer drugs (DREADD)-based approach to acutely enhance GABAergic neuronal transmission ^34^. Specifically, we injected an AAV mixture of synapsin-jRGECO1a and Dlx5/6-hM3Dq at P1-P2 and installed cranial windows at P8-P9 (Fig. 5A). We confirmed that the majority of jRGECO1a-expressing neurons were glutamatergic neurons, while Dlx5/6-hM3Dq-positive neurons were predominantly GABAergic (Fig. S5), as previously reported ^34^.

**Fig. 5.**
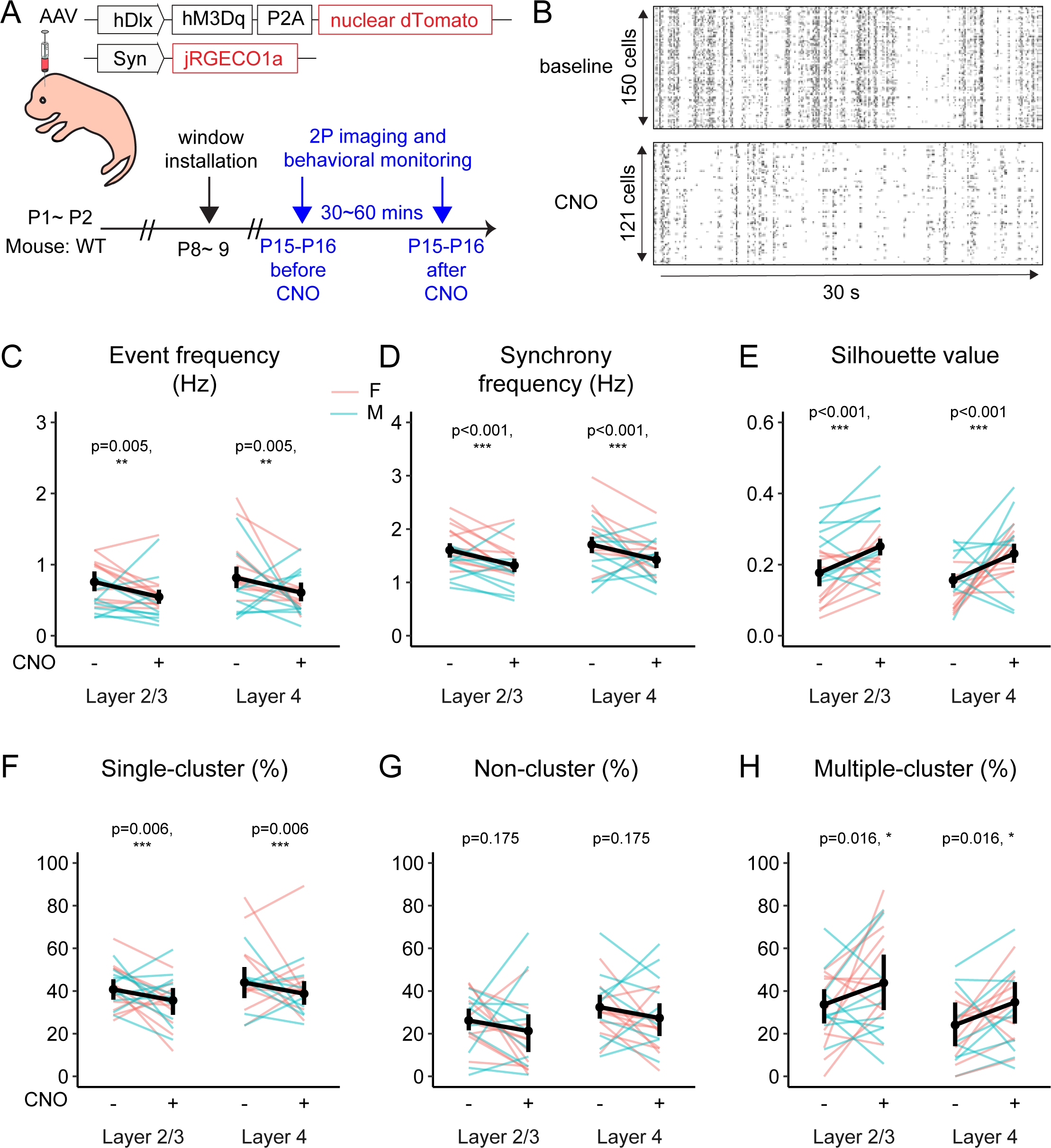
Enhancing GABAergic transmission reduced glutamatergic neuronal activity and enhanced subnetwork coherence. (A) The diagram illustrates the experimental design. Mice were co-injected with AAV-carrying jRGECO1a (AAV9-Syn.NES-jRGECO1a) and excitatory DREADD (AAV9-hDlx-hM3Dq-P2A-nuclear dTomato) at P1∼2. Following cranial window installation at P8∼9, pups were imaged at P15∼16. Imaging was performed both before and 30 minutes after the administration of CNO (1 mg/Kg). (B) Raster plot of spontaneous activity of L2/3 neurons before and after CNO administration. (C) Event frequency. (D) Synchrony frequency. (E) Silhouette value of event cluster. (F) Percentage of neurons participating in a single event cluster. (G) Percentage of neurons not involved in any event cluster. (H) Percentage of neurons participating in multiple clusters. Numbers of experimental mice: 6F, 4M. Data are presented as mean ± 95% confidence intervals derived from the bootstrapping output.

P15-P16, the midpoint of the developmental transition from P11-P21, was selected for assessing the impact of GABAergic inputs on neuronal subnetworks. After baseline calcium imaging, we administered the hM3Dq ligand, clozapine-N-oxide (CNO; 1 mg/kg, intraperitoneally), to enhance the excitability of GABAergic neurons (Fig. 5A). 30-60 minutes post-CNO treatment, calcium imaging was conducted to determine its impact. This acute enhancement of GABAergic activity significantly reduced the frequency of both individual calcium events (Fig. 5B, C) and synchronized events (Fig. 5D) in both L2/3 and L4 neurons and across both sexes. Despite the substantial reduction in event frequencies and synchronized events, CNO administration did not detectably impact the Pearson correlation coefficient or MI values of all neuronal pairs (Fig. S6A, B).

Results from subnetwork analysis showed that acute CNO treatment robustly increased silhouette values of event clusters in both L2/3 and L4, independent of sex (Fig. 5E), while the subnetwork numbers remained constant (Fig. S6C). The intensified tCC within the same subnetwork due to acute GABAergic enhancement was particularly evident in females in both L2/3 and L4 (Fig. S6D). Regarding the impact of GABAergic inputs on subnetwork configurations, we observed that CNO treatment decreased the percentage of single-cluster neurons (Fig. 5F) while increasing the percentage of multiple-cluster neurons (Fig. 5H). The percentage of non-cluster neurons was not significantly altered by CNO treatment (Fig. 5G). Analysis of the topographical distribution of neurons showed that CNO application had no detectable impact on the distances between neurons within the same subnetwork (Fig. S6E), the distance of individual neurons to their respective subnetwork centroid (Fig. S6F), or the pairwise distances between subnetwork centroids (Fig. S6G). Taken together, our data suggested that enhancing functional GABAergic inputs increases the sparseness of neuronal activity and cortical subnetwork cohesiveness without altering the anatomical distribution of the subnetworks.

## Discussion

In this study, we examined the development of S1 cortical subnetworks by imaging neuronal activity in both female and male mouse pups from P11 to P21, with a specific focus on examining periods of behavioral quiescence. Through the application of multiple clustering algorithms and rigorous statistical methods, including multilevel analysis with bootstrapping, we ensured the robustness of our findings. Our results reveal that cortical subnetworks are already present by P11, the earliest time point we assessed, and the number of subnetworks remains relatively stable from P11 to P21. However, these nascent networks undergo significant developmental refinement, marked by increased cohesiveness within individual subnetworks and greater distinction among the activity patterns of different subnetworks. Furthermore, our study with the DREADD approach provided compelling evidence that enhancing GABAergic inputs augments subnetwork coherence.

Understanding when and how neuronal subnetworks are established has been challenging, as training neonatal mice for specific tasks is impractical. A second challenge is the longitudinal imaging of developing pups that progress from minimal motor movements to abrupt increases in physical activity after eye-opening, such as walking and grooming. To overcome this challenge, we focused on analyzing spontaneous neuron activity patterns during periods of quiescence, when neurons within the same subnetwork often coactivate during spontaneous activity ^5,8,10–12^. By focusing on the behaviorally quiescent state, we identified three key changes of spontaneous activity from P11-P21: sparsification, desynchronization (characterized by burst activation of all neurons), and decorrelation (involving pairwise correlated activity of activity traces). These developmental changes were evident in both L2/3 and L4 layers and in both male and female mice (Fig. 1, Fig. 4, Fig. S2F-G, Fig. S4), aligning with previous studies ^15,17–21^. This approach provides a more comprehensive characterization of the cortical layer- and developmental trajectory properties of spontaneous network activity. Computational models of developing neural networks have demonstrated that blocking GABAergic transmission increases synchronous activity^35^. A recent study also demonstrated that somatostatin interneurons modulate parvalbumin interneurons and thus reduce the synchronization of cortical activity during P7-P11^36^. In our study, we found that enhancing GABAergic neuronal activity at P15 acutely reduces event frequency and synchrony frequency (Fig. 5C, D). Together, our findings suggest that the maturation of GABAergic inputs plays a significant role in developmental sparsification and desynchronization.

During cortical development, early postnatal spontaneous neuronal activity exhibited diverse patterns that contributed to the construction and refinement of functional network^37,38^. For example, as early as P1, S1 L4 neurons display a patchwork-type activity pattern driven primarily by peripheral inputs and independent of whisker movements^15,17^. This activity is thought to play a critical role in establishing the somatotopic map of whisker inputs^15^. In contrast, the transition from highly synchronous activity patterns dominating the early postnatal stage, to a mature state, characterized by sparse but co-activated distinct neuronal subsets (known as ensembles or subnetworks), is thought to enhance the efficiency of information storage, retrieval, and processing^5,6^. To identify neuronal subnetworks, defined as clusters of neurons exhibiting recurrent activity patterns arising from spontaneous activity, we first utilized k-means clustering, an unsupervised algorithm. During the event clustering process, we found that the k-means algorithm grouped approximately 25% of events, with about 1% of events being involved in event-multiple clusters (Fig. S7A). The relatively low number of events clustered may partly reflect the inherently sparse nature of neuronal activity, which is a common challenge in neuroscience. To further test the robustness of our clustering results, we employed additional similarity matrices, including cosine similarity and Jaccard similarity, for k-means clustering (Fig. S7B-C). Cosine similarity is particularly effective with dense data, where many attributes are present, and values are non-zero, as it considers the vector orientation. In contrast, Jaccard similarity is better suited for sparse data, where attributes are mostly absent or binary, focusing on the overlap between sets. When comparing the results obtained using these additional similarity matrices, we observed that the covariance matrix grouped the highest number of events and consequently formed more subnetworks (Figure. S7A-C). This finding indicated that the covariance matrix provided the optimal clustering performance for our data.

To validate the robustness of k-means clustering results and reduce the risk of random clustering effects, we also employed semi-supervised Louvain community algorithms and the DBSCAN algorithm. Louvain community detection is a greedy optimization method designed to extract non-overlapping communities from large networks. We tested two Louvain community detection algorithms, the uniform model and the asymmetric model, using the covariance matrix. The uniform model assumes that the network is undirected and unweighted, treating all edges equally and focusing solely on the structure to detect communities ^39^. In contrast, the asymmetric model accounts for the direction and weight of edges, allowing for the separate handling of positive and negative correlations ^40^. Different from the Louvain community method, DBSCAN defines clusters as dense regions separated by regions of lower density^28^. This approach allows DBSCAN to discover clusters of arbitrary shape while recognizing outliers as noise, making it particularly effective for datasets with non-linear structures. Our results revealed that k-means clustering and Louvain community detection clustered a similar percentage of events (Fig. S7A, D-E). However, DBSCAN clustered only about 10% of events (Fig. S7F). Notably, DBSCAN produced event clusters with higher silhouette values than the k-means and Louvain methods (Fig. S7A, D-F) and trended toward a heterogenous number of event clusters (Fig. S8E). DBSCAN’s clustering principle in predefining a similarity threshold (epsilon) affords its strength in identifying tightly grouped data points. In contract, method like k-means clustering is better suited for partitioning data to minimize overall variance. Of note, the k-means clustering algorithm identified more neuronal subnetworks compared to the Louvain community and DBSCAN (Fig. S7A, D-F). However, excitingly, all clustering methods consistently revealed the following outcomes: (1) developmental increases in silhouette values; (2) the stability in subnetwork numbers across P11-P21 (Fig. S7); (3) reductions in the percentage of single-cluster neurons and increases in the percentage of non-cluster neurons (Fig. S8). Suggesting the clustering outcomes we acquired are robust and it is unlikely that the observed decrease in the percentage of neurons participating in subnetworks is caused by the sparseness of neuronal activity.

During our investigation, most imaged neurons (∼90%) were identified as glutamatergic neurons, while approximately 10% were categorized as GABAergic neurons (Fig. S5E). It is plausible that both excitatory and inhibitory neurons contributed to the formation of subnetworks observed in our study. A computational study suggested that networks dominated by excitatory interactions exhibited rapid assembly formation, but this often involved the recruitment of additional, non-perturbed neurons, leading to nonspecific activation ^41^. Conversely, networks with strong excitatory and inhibitory interactions formed more confined assemblies, limited to the perturbed neurons^41^. In line with these findings, we observed a transition in subnetwork configuration from a high to lower participation rate between the 2^nd^ to 3^rd^ postnatal week (Fig. 4F-H), which coincides with an increase in cortical inhibitory inputs ^24,25^. Furthermore, the finding that acutely enhancing GABAergic activity reduced the percentage of neurons participating in subnetworks (Fig. 5F) provides strong support for the crucial role of inhibitory inputs in excluding neurons from existing subnetworks. Mechanisms involved in synaptic plasticity and maintaining excitatory/inhibitory balance are critical for subnetwork refinement. The majority of cortical glutamatergic neurons undergo developmental cell death during the first postnatal week, process that is mainly complete by P10^42,43^. Thus, it is unlikely that the observed decline in neuronal participation is due to this loss of excitatory neurons. On the other hand, the maturation of inhibitory synapses is influenced by several activity-dependent processes^44^. Thus, it is plausible that inhibitory circuit plasticity/refinement shapes subnetwork dynamics from P11-P21.

Following these results, an important question arises: what advantages do neurons gain from being restricted to individual subnetworks during development? In the mature S1 cortex, strong recurrent excitatory connectivity provides a robust framework for performing complex computational tasks ^45–48^. However, networks with high excitatory gain can be prone to instability. In contrast, inhibitory inputs provide a dynamic and flexible mechanism for stabilizing these potentially unstable networks ^48–50^. At P11, we noted that nascent event clusters exhibited the lowest silhouette values (Fig. 2D, 4D), likely reflecting an early stage when excitatory intra-cortical connections were just established ^26,27^ and inhibitory inputs remained weak ^24,25^. By P21, these event clusters showed high silhouette values, indicating either a more uniform activity pattern within individual subnetworks or more distinct patterns among different subnetworks. Importantly, this increase in similarity was observed when GABAergic neuronal activity was acutely elevated (Fig. 4E). These findings reinforce the idea that GABAergic inputs contribute significantly to enhancing the functional coherence of subnetworks during development. Mòdol et al. recently demonstrated that the number of neurons recruited into synchronous events decreases from P7/P8 to P11/P12, with functional assemblies initially restricted within cortical columns at P7/P8 and becoming more spatially dispersed by P11/P12^36^. Results from our imaging data from P11-P21 reveal a consistent number of neuronal subnetworks starting at P11. Taken together, we propose that cortical subnetworks are already formed by P7 and then undergo functional refinement until the 3^rd^ postnatal week as they acquire activity coherence within the same subnetworks, while remaining distinct from other subnetworks. In the visual cortex, the maturation of GABAergic inputs plays a pivotal role in shaping critical periods by regulating the timing and extent of plasticity^51–53^. Similarly, our study demonstrated that the refinement of neuronal subnetworks in the S1 cortex coincides with the maturation of GABAergic inputs^24,25^. This temporal alignment suggests that the developmental trajectory of subnetwork in the S1 cortex may be shaped by critical period plasticity. Such refinement likely reflects the influence of sensory input and environmental stimuli, which drive the fine-tuning of functional subnetworks to enhance cortical computation and integration during early development. Sensorimotor developmental studies reveal that mice begin bilateral whisking at P10 and actively explore their environment with their whiskers by P11^23,54^. It remains to be investigated how sensorimotor experience influences the assembly of neuronal subnetworks.

At P15, we observed that male mice exhibited larger silhouette values compared to female mice in both L2/3 and L4 (Fig. 4D). Our anatomical studies quantifying the densities of overall GABAergic input in the neuropil and perisomatic regions did not reveal any sex-dependent differences (Fig. S9A, C-E). Astrocytes interact with neurons via their delicate astrocytic end feet, which extensively develop from P7-P21 in S1^55,56^. Interestingly, astrocyte-specific translatomic analysis showed that the molecular signature levels matured faster in males than in females ^57^. In the hippocampus, studies suggest that increasing astrocytic calcium dynamics result in long-lasting synaptic potentiation ^58,59^. Future studies should be conducted to determine if astrocytes contribute to the sex differences observed in subnetwork properties.

Our anatomical studies quantifying the densities of overall GABAergic input identified differences between L2/3 and L4. Specifically, (1) L4 exhibited a higher density of putative parvalbumin synapses labeled by synaptotagmin-2 (Syt2) antibody ^60^, but not in total GABAergic synapses marked by vesicular GABA transporter (VGAT), compared to L2/3 (Fig. S9D-E). (2) L4 spiny stellate neurons not only have significantly smaller somatic volumes (Fig. S9F) but also receive fewer perisomatic GABAergic inputs than L2/3 neurons (Fig. S9G). Notably, L4 neurons within the same subnetwork displayed significantly lower tCC at P15 compared to L2/3 neurons (Fig. 4E). These findings suggest that parvalbumin inhibitory inputs may regulate correlated activity within the same subnetwork. Previous studies have shown that optogenetic or electrical stimulation of L2/3 pyramidal neurons in brain slices from the adult mouse visual cortex induced neuronal ensemble formation, accompanied by an increase in intrinsic excitability ^61^. Reducing this heightened intrinsic excitability reverses ensemble formation, highlighting its role in the process ^61^. Thus, future studies should investigate whether increasing intrinsic excitability in developing cortical neurons can facilitate the functional assembly of cortical subnetworks.

In summary, our study demonstrates early formation of subnetworks structures among S1 neurons in upper cortical layers, with a gradual increase in coherence during the 2^nd^ to 3^rd^ postnatal weeks. This maturation process involves retaining neurons with similar activity patterns within the subnetwork to enhance functional coherence. Furthermore, our findings highlight the role of GABAergic inputs in shaping subnetwork coherence.

## Resource Availability

Lead contact

Request for further information and resources should be directed to the lead contact Dr. Jui-Yen Huang

## Materials availability

This study did not generate new materials. Data and code availability

All raw data reported in this paper have been uploaded to DataJoint company (https://www.datajoint.com/) and are available upon requests from the lead contact. The data analysis workflow has been integrated into DataJoint elements for public access.

All codes for data analysis, statistical comparison test, and post-analyzed files can be accessed at https://github.com/EdnaHuang/SubnetworkAssembley2024”. Any additional information required to reanalyzed the data presented in this paper is also available from the lead contact upon request.

## Limitations of Study

We observed a reconfiguration of nascent subnetworks, characterized by a decrease in the percentage of neurons participating in each subnetwork (Fig. 3A-C, Fig. S4F-H). This transition from a larger group of neurons to a smaller, more selective neuronal pool may be essential for refining the activity coherence of nascent subnetworks, ultimately stabilizing their functional performance. Ideally, tracking the same groups of neurons over time would provide deeper insights. However, technical challenges related to brain expansion currently limit our ability to reliably trace more than 50 neurons for consistent clustering. Future studies could overcome this limitation by using cellular markers to identify and track the same neurons in longitudinal imaging studies. Additionally, our study provides the first experimental evidence demonstrating the role of inhibitory GABAergic inputs during the 2^nd^ and 3^rd^ postnatal weeks in neuronal subnetwork assembly. Further investigation is needed to elucidate the precise mechanisms, timing, and specific GABAergic subtypes in refining neuronal subnetworks. For example, inhibiting the activity of a particular subtype of GABAergic neurons during specific development ages by combining Cre-lines with advanced genetic manipulation approaches^62^ to determine the acute and long-lasting impacts of GABA on subnetwork properties. Such knowledge will allow us to acquire a more in-depth understanding of the sensorimotor integration deficits that occur in various developmental neurological disorders and sex-dependent vulnerabilities.

## Acknowledgments

We thank Bruna Baumgarten Krebs, Joseph Kimmel Leffel, and Sarah Van Meter for their technical support, and Drs. Carlos Portera-Cailliau, Duton Oyesanmi, Ju Lu, Nathalie L. Rochefort, Ruth Eberle, and Sander W. Keemink for their coding assistance. We are grateful to clinical veterinarian Randalyn Shepherd for her support and valuable suggestions to maintain the health of post-surgical pups. We also thank Drs. Ehren Lee Newman, Richard Betzel, and Zoran Tiganj for their critical feedback on additional clustering analyses. We appreciate the assistance of Eric Wernert, Esen Tuna, and James Robert McCombs with data storage and high-performance computing resources. We thank Miss Pei-Ying Chen from the Indiana Statistical Consulting Center for her support with statistical analysis, and Drs. Cary Lai, Ehren Lee Newman, Ken Mackie, Richard Betzel, Sen Yang, Wei-Hsiang Huang, and Zoran Tiganj for their helpful comments.

This work was supported by the following grants: National Institutes of Health grant R01NS086794 (HCL), National Institutes of Health grant P30DA056410 (HCL), startup funds from the IUB College of Arts and Sciences (HCL), and Gill endowment funds (HCL). Additional support came from the Indiana Clinical and Translational Sciences Institute, funded in part by Grant Number UL1TR002529 from the National Institutes of Health National Center for Advancing Translational Sciences, Clinical and Translational Sciences Award. The content is solely the responsibility of the authors and does not necessarily represent the official views of the National Institutes of Health. We also acknowledge support from Lilly Endowment, Inc., through its contributions to the Indiana University Pervasive Technology Institute, and Shared University Research grants from IBM, Inc., to Indiana University. This work utilized the Extreme Science and Engineering Discovery Environment (XSEDE), supported by National Science Foundation grant number ACI-1663578.

## Author Contributions

Conceptualization: HCL, JYH

Methodology: HCL, JYH, MH

Software: AB, JYH, MH, SJB, XL

Validation: AB, JYH, MH

Formal Analysis: AB, AJX, JYH, MH

Investigation: AJX, JYH, LNH, XL

Resources: HCL

Data Curation: AJX, JYH, MH, SJB, XL

Writing—original draft: JYH, MH, HCL

Writing—review & editing: AB, AJX, HCL, JYH, LNH, MH, SJB, XL

Visualization: HCL, JYH, MH

Supervision: HCL, JYH

Project Administration: JYH

Funding Acquisition: HCL

## Declaration of Interests

The authors declare no competing interests.

## STAR Methods

### Key Resources Table

### Chemicals and Antibodies

| REAGENT or RESOURCE | SOURCE | IDENTIFIER |
| --- | --- | --- |
| **Antibodies** |  |  |
| Mouse anti-CamKII | AbCam | Cat# ab22609; RRID: AB_447192 |
| Chicken anti-GFP | Aves | Cat# GFP-1010; RRID: AB_2307313 |
| Rabbit-anti RFP | Rockland Antibodies | Cat# 600-401-379; RRID: AB_2209751 |
| Goat anti-chicken Alexa Fluor 488 | Thermo Fisher Scientific | Cat# A11039; RRID: AB_2534096 |
| Goat anti-rabbit Alexa Fluor 555 | Thermo Fisher Scientific | Cat# A21429; RRID: AB_2535850 |
| Goat anti-mouse Alexa Fluor 647 | Thermo Fisher Scientific | Cat# A21236; RRID: AB_2535805 |
| **Bacterial and virus strains** |  |  |
| pAAV9. Syn.NES-JRGECO1a | <sup>63</sup> | RRID: Addgene_100854 |
| pAAV9.hDlx-GqDREADD-dTomato | <sup>34</sup> | RRID: Addgene_83897 |
| **Chemicals, peptides, and recombinant proteins** |  |  |
| KCl | Sigma-Aldrich | Cat# P9541 |
| Trizma-HCl | Sigma-Aldrich | Cat# T5941 |
| Triton X-100 | Sigma-Aldrich | Cat# X-100 |
| Proteinase K | Roche | Cat# 3115828001 |
| EconoTaq Plus Green 2x mater mix | Lucigen | Cat# 30033-1 |
| FastGreen FCF dye | Sigma-Aldrich | Cat# F7252 |
| Clozapine-n-oxide | Sigma-Aldrich | Cat# C0832 |
| Paraformaldehyde | Sigma-Aldrich | Cat# 8.18715 |
| Ethylene glycol | Sigma-Aldrich | Cat# 324558 |
| polyvinyl pyrrolidone | Sigma-Aldrich | Cat# PVP360 |
| Normal goat serum | Jackson ImmunoResearch | Cat# 005-00-121 |
| DAPI | Thermo Fisher | Cat# D3571 |
| Prolong Diamond Antifade mounting medium | ThermoFisher | Cat# P36961 |
| **Experimental models: Organisms/strains** |  |  |
| Wildtype C57/B6 mice | Jackson Laboratory |  |
| R26-LSL-hM4Di-DREADD | Jackson Laboratory | RRID: IMSR_JAX:026219 |
| R26-LSL-Gq-DREADD mice | Jackson Laboratory | RRID: IMSR_JAX:026220 |
| FGF9 flox | 64 | RRID: IMSR_JAX:030675 |
| Nex-Cre | 65 |  |
| GAD67-GFP | 66 |  |
| Software and algorithms |  |  |
| NIS Element | NIKON microscope | https://www.microscope.healthcare.nikon.com/ |
| Imaris | OXFORD instruments | https://imaris.oxinst.com/ |
| Python 3 | Python Software Foundation | https://www.python.org/ |
| Adobe Illustrator | Adobe | https://www.adobe.com/ca/products/illustrator.html |
| MATLAB2023 | Mathworks | https://www.mathworks.com/products/matlab.html |
| FIJI | 67 | https://imagej.net/software/fiji/ |
| Suite2p | 68 | http://www.suite2p.org |
| FISSA | 69 |  |
| DeepLabCut | 70 |  |
| Facemap | 71 |  |
| R4.2.2 |  | https://www.r-project.org/ |
| RStudio |  | http://www.rstudio.com/ |

#### Experimental Design

Most of the animals used in this study were of a C57/B6J background. To increase the sample size for meaningful statistical comparisons, we also included animals from genetically modified lines, which, although primarily C57/B6J, were less pure. In figures 1 and 2, Some animals in figures 1 and 2 are mixed C57/B6J with R26-LSL-hM4Di-DREADD or R26-LSL-Gq-DREADD or FGF9 cKO mice (NEX-Cre negative, *FGF9^flox/flox^*, GAD-GFP). All animals in figure 5 were C57/B6J mice. Both male and female mice were used for data analysis. All procedures involving animals were performed in accordance with the Indiana University Bloomington Animal Research Committee, and the NIH Guide for the Care and Use of Laboratory Animals.

### METHOD DETAILS

#### P1/P2 injection of AAV for jRGECO1a expression

The surgical procedure was conducted as described previously ^18^ with slight modification. pAAV9. Syn.NES-JRGECO1a (100854-AAV9) and pAAV9.hDlx-GqDREADD-dTomato (83897-AAV9) viruses were purchased from Addgene and diluted to a working titer of 1x10^13^ GC/ml with 0.05% FastGreen FCF dye. Postnatal one- or two-day pups were anesthetized with isoflurane (5% for induction, 2-3% maintenance via a nose cone, vol/vol). Subcutaneous injection of ketofen (5mg/kg) was administered. The surgical region was disinfected with 4% chlorhexidine and 70% alcohol. We used soft silicone ear bars to position the mouse on the stereotaxic frame to make the desired injection area (presumptive somatosensory cortex) as flat as possible. We used scissors to create a 3-4 mm triangular skin flap over S1. The periosteum was gently cleared under the skin flap using a sterile cotton swab. The skin flap (folded back) was covered with a piece of saline-soaked Gelfoam to prevent the skin from drying during the procedure. At the injection site, the bone was drilled lightly to create a small crack, permitting injection via pulled-glass capillary without exposing the dura. Glass micropipettes (O.D. 1 mm, I.D. 0.5 mm Sutter Instruments, cat no. BF100-50-10) pulled by using a P1000 pipette puller (Shutter Instruments, CA, USA) were used to inject ∼200 nL of AAV into the superficial cortex, using an injection pump equipped with a Hamilton syringe. The pipettes were positioned for injection at 70∼90° angle to the skull surface and gently pierced through the cracked bone and into the superficial cortex. The injection flow rate was 50 nL/min. The light blue dye was visible on the brain surface of pups with successful injection. The glass pipettes were left in place for one minute before withdrawal. The skin flap was sealed with a small drop of VetBond to the injection site. The entire surgery was complete in 10-15 min per pup. The pups recovered on a warm blanket before being returned to the dam.

#### Cranial window surgery

The surgical procedure was conducted as previously described ^18^ with slight modification. Mice aged at P8∼ 9 were anesthetized with 5% isoflurane for induction, followed by 2-3% maintenance (vol/vol, via nose cone). Pups also received Ketoprofen (5 mg/kg, s.c.), Dexamethasone (1 mg/kg, s.c.), and a mixture of lidocaine and epinephrine (4 mg/kg and 0.1 mg/kg, respectively, applied topical to the surgical region) for analgesia and anti-inflammatory purposes. Pups were positioned using blunt ear bars on the stereotaxic frame. The original AAV injection site was visible as a well-healed triangular scar on the skin. The pup’s head was swabbed with 4% chlorhexidine and 70% alcohol to disinfect the skin and the skin over the surgical area was removed. A dental drill was used to carve a circular craniotomy, 3 mm in diameter. The drilling was stopped once the bone had cracked around the perimeter of the craniotomy. Any blood was promptly cleared to prevent a blurred window. The entire drilled area was soaked with sterile saline using Gelfoam for at least 3 min to soften the bone. The bone fragment was gently removed, and a 3 mm glass plug coverslip (generated by adhering 2.3 mm coverslip with optical transparent UV-activated adhesive) was positioned over the craniotomy. The glass was sealed using cyanoacrylate then dental acrylic (Metabond). A custom-made circular head plate (less than 0.5 g, Fig. S1C) was positioned over the area, centered around the cranial window, and was adhered to the skull using dental acrylic (Metabond). The pup was then placed on a heating pad with bedding for recovery and then returned to the dam when fully mobile. Post-surgical pups were fed with kitten milk replacement (every 4-5 hours during daytime) to provide nutrition if pups did not gain body weight and had no milk spot in the belly. Pups became more active in seeking the dam for milk two days after the surgical procedure. Pups were subjected to habituation on the imaging station for 15 minutes the day after surgery. Following that, pups were habituated on the imaging station for 30 minutes (the average imaging time) or subjected to the imaging procedure daily. During our imaging period, pups continued to gain body weight as they grew up (Fig. S1E).

#### In vivo calcium imaging in head-restrained neonatal pup

Before imaging, a metal bar was attached to the headframe to provide the anchoring point to the imaging station (Fig. S1B, C). Calcium imaging was performed on a Nikon A1R MP^+^ two photon microscope, with an Insight DeepSee pulsed laser (Spectra-Physics Inc, Santa Clara, USA), resonant scanner, a 25x, 1.1 NA water immersion lens (Nikon, Japan), and Nikon Elements software. The homemade imaging station was equipped with a lightweight foam disc covered by a circular paper drape. The center of the disc was fixed, allowing pups to rest, whisk, or walk with minimal friction (Fig. S1D). A heating pad was placed under the imaging platform to maintain the body temperature of neonatal pups. The light source was an Insight laser at 1040 nm for jRGECO1a. Whole field images were acquired at 65.6 ms/frame (∼15.23 Hz) with a resolution of 512 x 512 pixels (representative Videos in Videos S1, S2, and S3). The pups’ body movements were simultaneously monitored and recorded during the imaging procedure using infrared cameras (Fig. S1D). USB cameras (30 Hz) with an 850 nm IR LED light source and IR-cut filter at 850 nm were used to record body movements. A Flea3 USB3 camera (100 Hz) with an IR-cut filter at 850 nm was used to record whisker movements. Before calcium imaging, pups were habituated on the imaging station for at least 10 mins. To find the same field of view to image in subsequent imaging sessions, we used the vasculature pattern as a landmark to locate the previous imaging position (Fig. S1F). First, the global vasculature pattern in the cranial window was imaged by using a dissecting microscope (Fig. S1F upper panel). The location where calcium imaging was performed was marked on the map. Next, the pial surface vasculature pattern was imaged under the two-photon microscope and used as a second landmark (Fig. S1F lower panel). Each imaging session was conducted within one hour per animal to minimize the adverse effects of maternal separation. For imaging in L2/3, the location was set at 120-150 µm below the pia surface, while for L4, it was adjusted to 200-350 µm below the pia surface, following the guidelines provided by Van der Bourg et al. (2017)^23^. After the final imaging session, some pups were perfused to confirm the imaged area was in S1 (Fig. S1G). Post hoc analysis confirmed that the imaged area spanned from bregma -0.67 mm to -1.84 mm encompassing regions such as the S1 barrel field, shoulder region, dysgranular zone, and trunk region.

#### Data analysis for calcium imaging

The preprocessing of all raw calcium movie data was done using a Suites2P toolbox ^68,71^. Briefly, Suite2p first aligned all frames of a calcium movie using two-dimensional rigid registration based on regularized phase correlation, subpixel interpolation, and kriging. Suite2p computed a low-dimensional decomposition of the data using a clustering algorithm that finds regions of interest (ROIs) based on the correlation of the pixels inside them. We inspected every ROI using the following criteria: (1) the classical morphology should be a donut-shaped circle because the cytosolic distributions of jRGECO1a; (2) the neurons with obvious nuclear signals were excluded, as were cells from animals injected with pAAV9-hDlx-GqDREADD-dTomato expressed nuclear dTomato fluorescent protein (Fig. S5); (3) ROIs never overlapped. For the neuropil correction, we used the Fast Imaging Signal Separation Analysis (FISSA) toolbox ^69^. The coordinates of selected ROIs and post-motion-corrected tiff files were imported to the FISSA toolbox for neuropil decontamination. After neuropil correction, the ΔF/F was calculated as (F-F0)/F0. Where the F is the extracted fluorescence signal, and F0 is the median of the bottom 10^th^ percentile of the fluorescent trace. To normalize the differential fluorescent change caused by the expressed jRGECO1a level in each neuron, we further calculated the modified Z score for each neuron ^72,73^. Zscore = F(t) - mean(baseline) / std (baseline). Where the baseline period is the 10-s period with the lowest variation (s.t.d.) in ΔF/F. We used the Peakfinder function in MatLab to find the conceivable calcium transient referred to neuronal spike activity (Fig. S1D). The detected calcium event is defined as the fluorescent peak amplitude greater than four times the noise level. The noise level is the mean Z score within five second-time windows when it has the lowest root mean square (Fig. S1D). To analyze the spontaneous neuronal activity, we used the DeepLabCut toolbox ^70,74^ to annotate the movement of the limbs and identify the time windows when pups show minimum activity (stationary condition). In addition, the video-recorded facial movements were analyzed by Facemap Python package ^71^. Both annotations were plotted to visualize the overall movement of the body and whisker (Fig. S1E). For spontaneous activity quantification, the time period with minimum body movement and whisking was used for calculating various parameters (Fig. S1E).

The synchrony frequency analysis was performed as described previously ^20^. We reconstructed activity histograms that plotted the percent of active cells for each frame (Fig. 1E). The threshold for detecting a network event was set by repeatedly shuffling each deconvolved somatic calcium trace 100 times while maintaining the average firing rate for each cell. A surrogate activity histogram was constructed from each reshuffled trial. The threshold was calculated by first finding the peak percentage of cells active in each reshuffled trial, and then sorting these values to find the 95^th^ percentile, which was set as the threshold for significance (p=0.05). The time point when the percent of active neurons exceeded this threshold was set as the start of an event. The time point when the percentage of active neurons fell below this level was set as the end of the event. The mean frequency of peaks of synchrony and the mean fraction of active cells at the maximum point of each peak of synchrony were calculated. For pairwise correlation analysis, Pearson correlation coefficients and mutual information were computed using Z-score normalized traces acquired during quiescent states. Additionally, we performed tCC analysis on deconvolved neuronal activity.

To identify the neuronal subnetwork, we employed a similar strategy as previously described ^11,16^. The initial calcium event clusters were identified using a clustering algorithm based on calcium dynamic similarity. Unsupervised clustering of calcium events was obtained by using the k-means algorithm on the normalized covariance matrix (frame-frame) with cluster numbers ranging from 2-19. One hundred iterations of k-means were run for each cluster number, and the iteration that resulted in the best-average silhouette value was retained. Each cluster was then associated to a cell assembly that comprised those neurons that significantly participated in the events within that particular cluster. Cell participation in a given cluster was considered statistically significant if the fraction of synchronous events in that cluster that activated the cell exceeded the 95^th^ percentile of reshuffled data. If a neuron was significantly active in more than one cluster, it was associated with the one in which it participated the most (based on percentage). The overlap between assemblies/subnetworks was quantified by calculating the silhouette value of each cell (with the normalized hamming distance between each cell pair as a dissimilarity metric). A neuron was significantly involved in a single cluster if its silhouette value was higher than expected by chance (95^th^ percentile after reshuffling). Events were finally sorted with respect to their projection onto neuronal subnetworks (Fig. S3A-B). Alternatively, neurons were grouped according to their subnetwork identities (Fig. 2C). We computed the tCC for detected event (binary format) for all possible neuronal pairs or neuronal pairs within their subnetworks or between subnetworks.

To evaluate the robustness of the k-means clustering results, we compared our results to those obtained from two alternative similarity metrices: cosine similarity and Jaccard similarity. For each alternative metric, we replaced the covariance matrix with the respective alternate values and completed the remaining steps of analyses identically. In addition to unsupervised k-means clustering algorithm, we also utilized semi-supervised Louvain community detection algorithm ^75–77^ and DBSCAN^28^. Two commonly used models of Louvain community detection algorithm, the uniform null model ^39^ and the asymmetric model ^40^, both developed for the neuroscience field, were chosen to cluster the covariance similarity matrix. These models were applied to the covariance similarity matrix, running 500 iterations for the uniform models and 100 iterations for asymmetric models, followed by 10 iterations of consensus clustering without removing weak elements ^78^. The iteration numbers were chosen based on clustering result reach converge. For DBSCAN, we converted the covariance similarity matrix (CovM) into a distance matrix using the formula distMatrix = 1 - CovM. DBSCAN clustering was then performed using a maximum distance threshold (epsilon) and a minimum number of points (minPts) to form a cluster. The optimal epsilon value for each dataset was identified using K-distance graphs^79^ and minPts was set to 3 to ensure inclusion of most data points as core points. Once event clusters were identified by either Louvain community detection or DBSCAN, the remaining analysis steps were performed in the same manner as for k-means clustering to maintain consistency across methods.

#### Immunohistochemistry, imaging, and quantification

Mice were deeply anesthetized with isoflurane and subsequently perfused with 4% paraformaldehyde in phosphate-buffered saline (PBS) at a rate of approximately 2 mL/minute for a duration of ten minutes. Fixed brains were then sectioned into 100 μm sections in the coronal plane using a Leica VT-1000 vibrating microtome (Leica Microsystems) and stored in an antigen-preserved solution (consisting of PBS, 50% ethylene glycol, and 1% polyvinyl pyrrolidone) at -20 °C until further analysis. Sections were permeabilization with 2% Triton X-100 and then incubated in a blocking solution (3% normal goat serum in PBS with 0.3% Triton X-100). Primary antibodies, prepared in the blocking solution, were applied overnight. Detection of primary antibodies was achieved using the appropriate secondary antibody conjugated with an Alexa series fluorophore was utilized. DAPI (100 ng/ml, ThermoFisher) was included in the secondary antibody solution to stain nuclei. For cell counting, Z-stack confocal images were captured from both hemispheres using a Nikon A1 confocal microscope equipped with a 10X/NA0.45 objective. Z-stacks were acquired at 1 µm intervals, resulting in a total imaged thickness of 5 µm. Quantification of colocalization between different markers, was performed using single-plane images, and analysis was carried out with NIH ImageJ software.

For the quantification of inhibitory synapses, Z-stack confocal images were obtained using a Nikon A1 confocal microscope with a 60X/NA1.4 objective at three times software zoom. The Z-stack was captured at 0.1 µm intervals, and each Z-stack contained 1 to 3 somas per image. Two images from each hemisphere were captured for each animal, and images from both hemispheres were analyzed using Imaris software (Bitplane, Zurich, Switzerland) following a workflow based on previous published methodologies ^80–85^. Given the variability in the volume occupied by nuclei and vasculature in each image, which influenced synaptic marker density quantification, surface objects of nuclei and vasculature-like structures were created using the Imaris surface module. The VGAT- or Syt2-channel was masked by these objects to exclude the volume they occupied. The post-masked VGAT channel was used to generate a surface object (referred to as the neuropil object) for analysis. To analyze VGAT and Syt2 double-positive puncta, the ImarisColoc function was employed to generate a new channel containing the VGAT and Syt2 colocalized signal, which was then used for subsequent spot detection. For spot detection, synaptic puncta labeled with VGAT, Syt2, and VGAT/Syt2 colocalized puncta were detected using the Imaris spot module with a diameter threshold of 0.5 μm. Synaptic puncta typically range in diameter ranging from 0.25–0.8 ^86,87^. To determine the optimal detection threshold, the threshold was manually defined for one image from each animal, and the threshold that detected the most synaptic puncta without creating artifacts was applied to analyze all images. This generated a spot layer for each synaptic marker channel. Synaptic density was calculated as the number of detected synapses inside the neuropil object divided by the neuropil volume. For peri-somatic inhibitory synapse quantification, the surface objects were created based on Kv2.1 signal using the surface module. VGAT, Syt2, and VGAT/Syt2 double-positive puncta located within 0.25 μm of the soma object were defined as perisomatic inhibitory synapses. All image acquisition and data analysis procedures were conducted in a blinded manner.

#### Statistical analysis

Given the clustered nature of the experimental design ^29–31^, we primarily employed linear mixed-effects (LME) models for multilevel statistical analysis to determine the statistical significance for all data. In cases where the data did not exhibit clustering, we fitted linear regression (LM) models instead to assess the statistical significance. We further refined the models by removing interaction terms that did not improve model fits based on measures such as Aikake Information Criterion and Bayesian Information Criterion values. To obtain reliable and unbiased estimates of variance components in both LME and LM analyses, we employed a bootstrapping procedure to resample the dataset (n=10,000) to create simulated samples ^29,30^. With this information, we conducted *post-hoc* comparisons using the bootstrapping output as the final statistical results. Each data point or line presented in the figures represents an individual dataset. Original data values and detailed statistical comparisons for all figures are available at https://github.com/EdnaHuang/SubnetworkAssembley2024. As the data were not normally distributed, we employed Spearman correlation coefficient to assess the correlation between subnetwork number and neuronal number or time intervals.

## Supplemental information

Document S1. Figures S1-S9

Videos S1. Calcium imaging from P11 pups. Post X-Y motion-corrected calcium imaging video of L2/3 neurons from P11 mouse pups. The video was acquired at 15 frames per second.

Videos S2. Calcium imaging from P15 pups. Post X-Y motion-corrected calcium imaging video of L2/3 neurons from P15 mouse pups. The video was acquired at 15 frames per second.

Videos S3. Calcium imaging from P21 pups. Post X-Y motion-corrected calcium imaging video of L2/3 neurons from P21 mouse pups. The video was acquired at 15 frames per second.

## Notes

https://github.com/EdnaHuang/SubnetworkAssembley2024

### Competing Interest Statement

The authors have declared no competing interest.

### Summary of Updates

Adding an additional clustering algorithm (DBSCAN) and including mutual information calculation for the correlation study. Revising the discussion section to incorporate the reviewer's comments.

