## Supplementary data for "From Initial Formation to Developmental Refinement: GABAergic Inputs Shape Neuronal Subnetworks in the Primary Somatosensory Cortex"

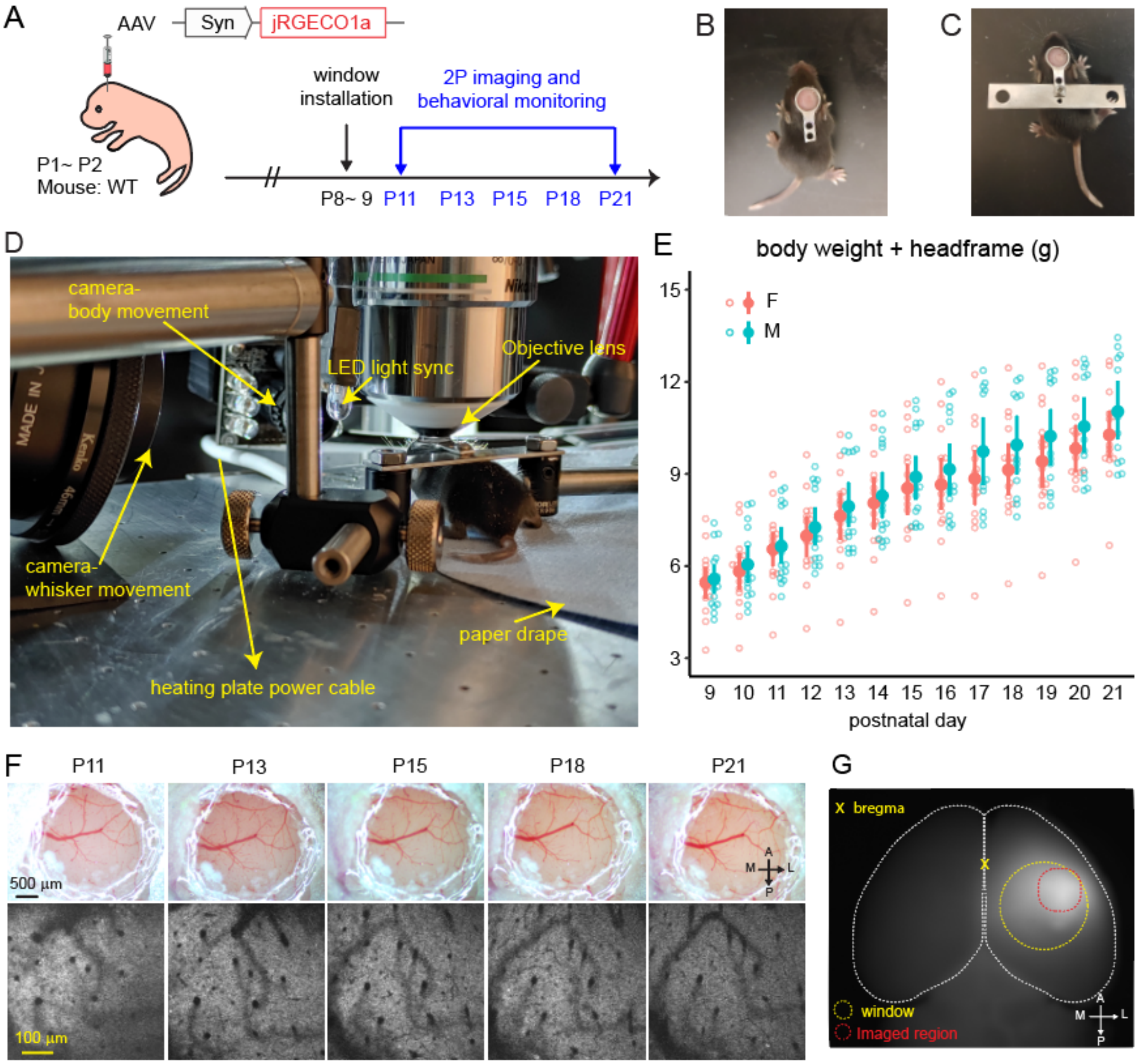


**Fig. S1. Setup and imaging of neonatal mice for calcium imaging studies**

(A) Schematic representation of the experimental timeline. (B) Attachment of a stainless-steel armature to the skull, surrounding the cranial window. (C) Metal bar used for providing anchoring points on the armature. (D) Photo of a P13 neonatal pup on the imaging station. A heating plate embedded under the metal platform keeps the pup warm. The center of a lightweight foam disk, covered by a circular piece of blue paper drape, is fixed. The disk rotates as the pup walks, allowing free limb movement while the head remains fixed for imaging. (E) Body weight changes of pups during the imaging period, with data presented as mean ± 95% confidence intervals of original data. Numbers of experimental mice. 15 females (F), 15 males

(M; 3 pups did not reach P21). (F) Upper panel: Example bright field images showing the cranial window implanted at P9, which remains optically transparent during the investigation. Lower panel: Example images showing the vasculature pattern under the two-photon microscope at different ages. (G) Dorsal view of brain tissue from a mouse with a window installed at P8 and perfused at P30. The imaging area is restricted to the upper right corner to target the somatosensory cortex. A, anterior; P, posterior; M, midline; L, lateral.

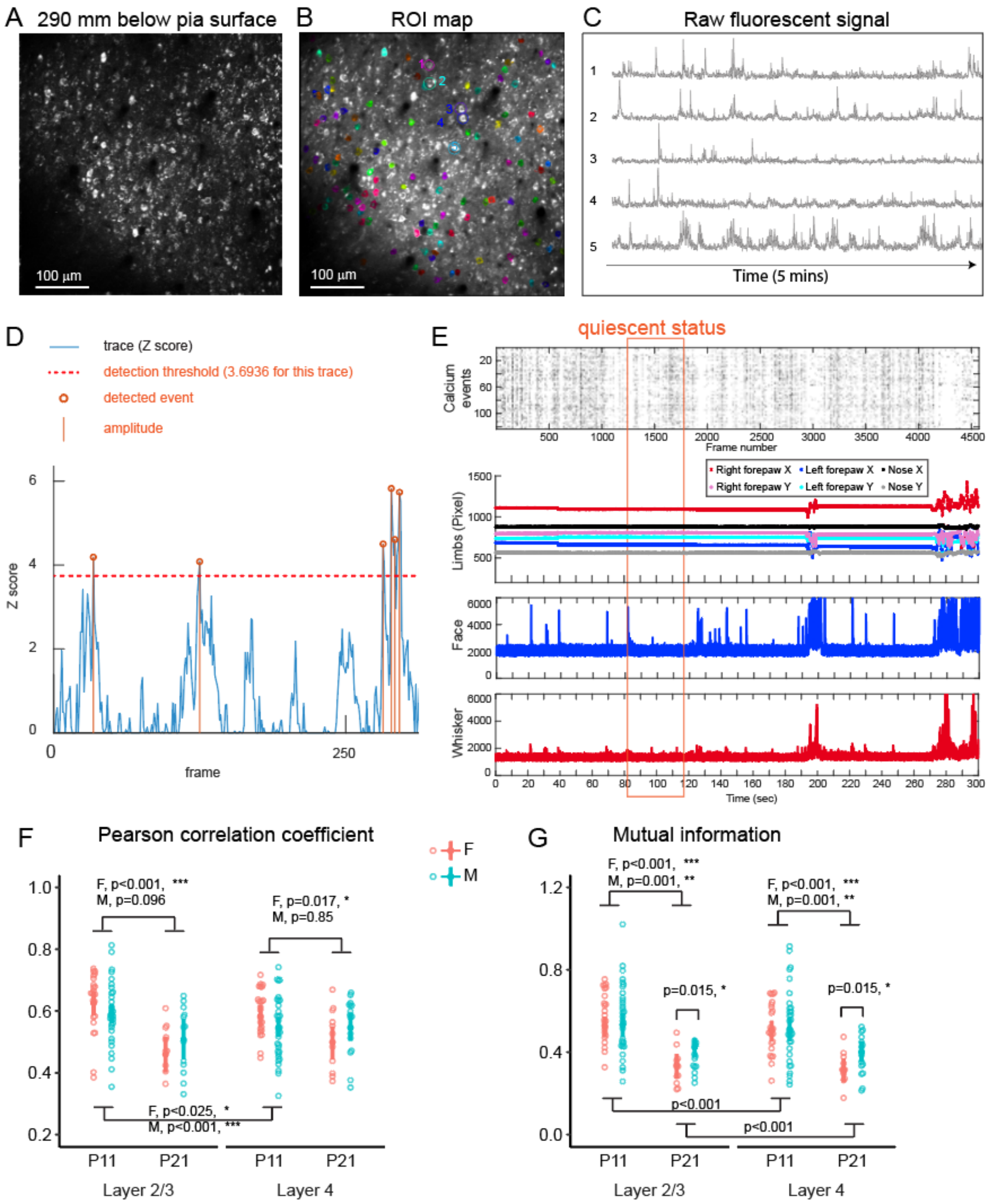


**Fig. S2. Calcium imaging data analysis workflow.**

(A) A representative calcium movie (maximum intensity projection across time) from a P15 mouse. (B) Region-of-interest (ROI) map after manual curation. (C) Exemplar calcium signal traces from 5 neurons in B. (D) Exemplar normalized Z-score fluorescent trace. Calcium transients above the detection threshold (four times the noise level, see Materials and Methods for details) are considered neuronal spike activity. (E) Exemplar figure showing detected calcium event, body movement, facial, and whisker movement. Alignment of neuronal activity with behavior allowed selection of analysis times during minimal body movement (quiescent status). (F) Pairwise Pearson correlation coefficients for all neuronal pairs. (G) Mutual information analysis for all neuronal pairs. Animal numbers are consistent with Fig. 1. Data are presented as mean ± 95% confidence intervals derived from the bootstrapping output.

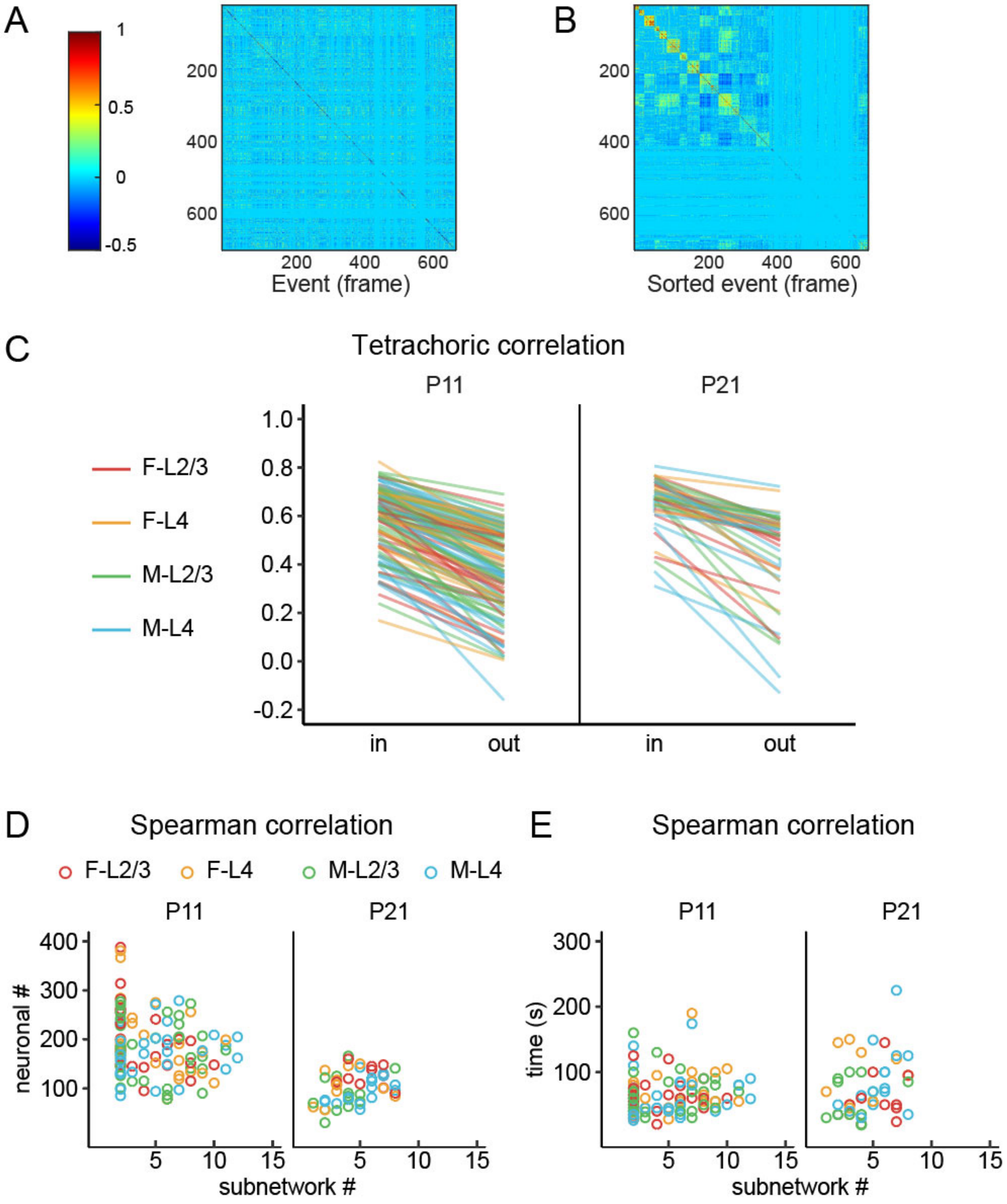


**Fig. S3. The spontaneous neuronal activity gradually de-correlated in both L2/3 and L4.** (A) The covariance similarity matrix generated by activity event (frame-frame) in the P11 dataset from Fig. 2A. (B) Re-sorted covariance similarity matrix based on an unsupervised k-means clustering algorithm, identifying seven event clusters in the dataset, with approximately 200 frames remaining unclustered. (C) Tetrachoric correlation coefficient (tCC) for neuronal pairs within the same subnetwork (in) and between different subnetworks (out). Notably, the tCC for neuronal pairs within the same subnetwork at P11 and P21 was consistently higher than for pairs between different subnetworks. (D) Correlation between neuronal numbers and subnetwork numbers. Spearman correlation coefficient, $\rho = -0.063$. (E) Correlation between time intervals included for clustering and subnetwork numbers. Spearman correlation coefficient, $\rho = 0.233$. Each line or circle represents one dataset.

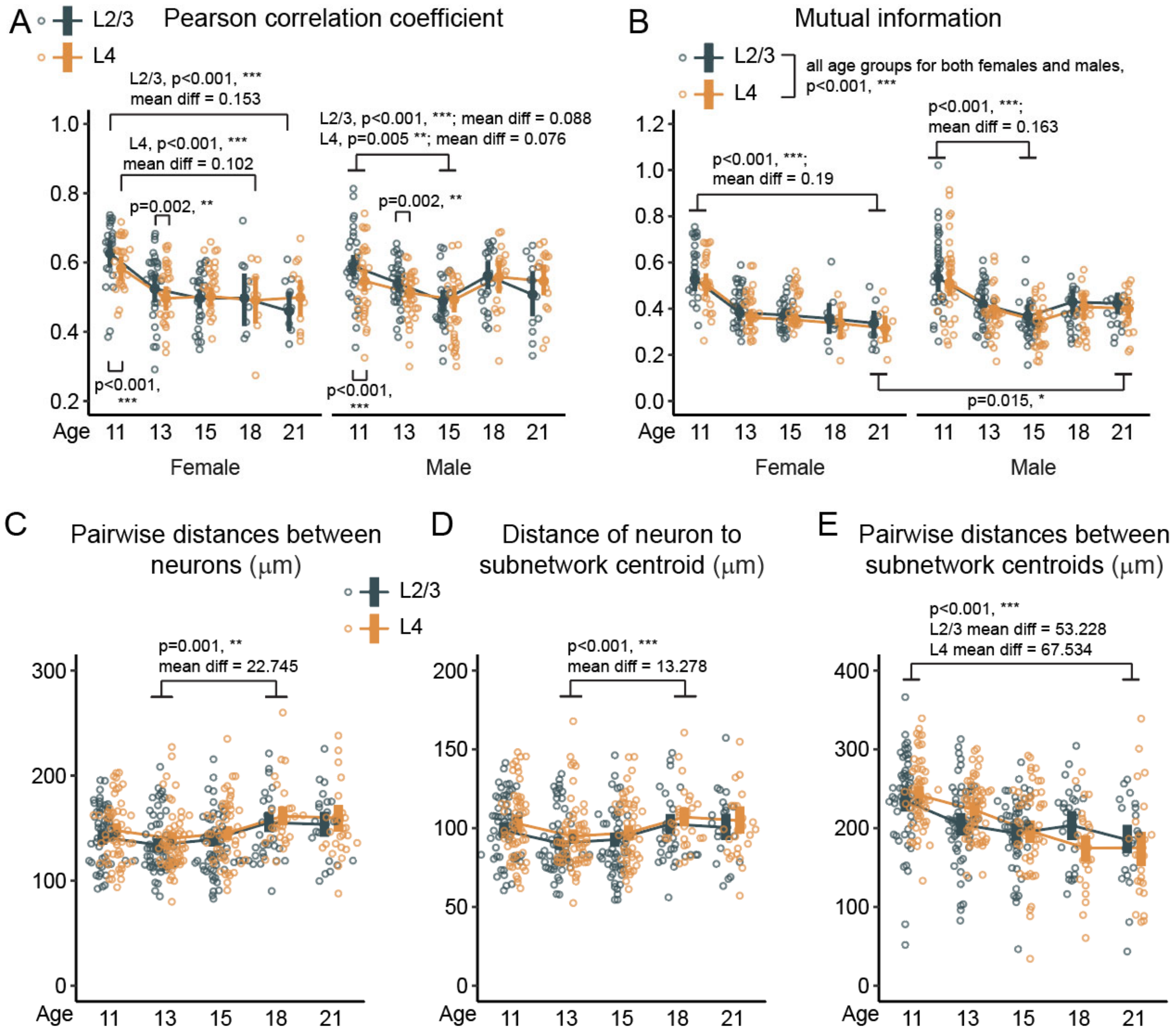


**Fig. S4. Developmental trajectory of cortical subnetwork assembly.**

This figure summarizes the progression of various parameters associated with cortical subnetwork development from P11 to P21. (A) Pearson correlation coefficient for all neuronal pairs. (B) Mutual information analysis for all neuronal pairs. (C) Pairwise distances among neurons within the same subnetwork. (D) Distances of neurons to their respective subnetwork centroids. (E) Pairwise distances between subnetwork centroids. Data from both sexes were combined in panels where no sex-dependent effects were observed to streamline the presentation. Animal numbers are consistent with Fig. 4. Data are presented as mean ± 95% confidence intervals derived from the bootstrapping output.

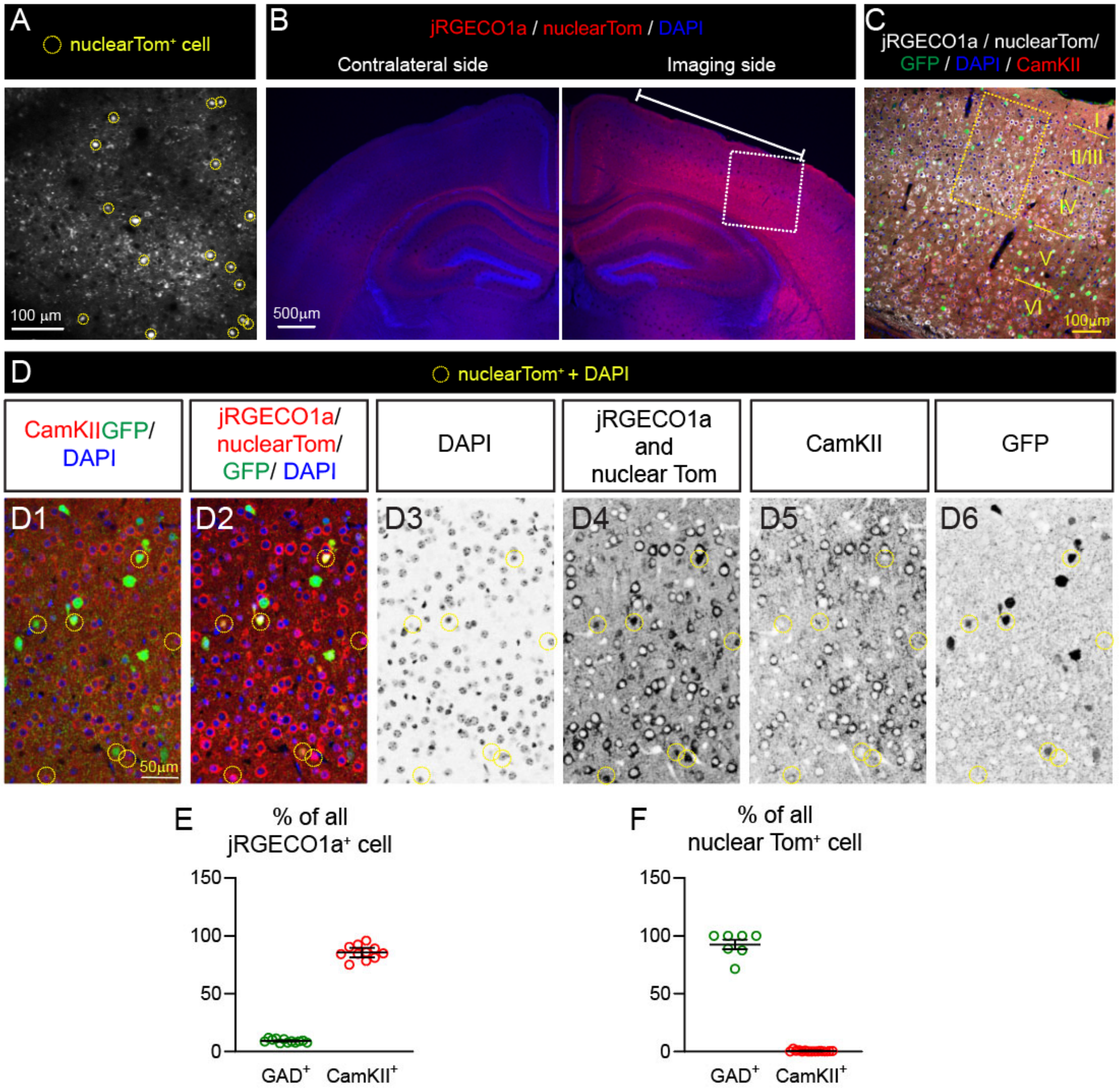


**Fig. S5**. **The majority of jRGECO1a-expressing neurons were glutamatergic neurons.** (A) Representative calcium movie (maximum intensity projection across time) from a P15 mouse injected with an AAV carrying excitatory DREADD. Yellow dashed circles indicate neurons expressing nuclear dTomato red fluorescent protein (Tom), which were excluded from data analysis during ROI curation. (B) Brain sections from a P21 mouse post-calcium imaging. The white line (2.3 mm) indicates the region where the cortical surface is slightly recessed due to the window placement. (C) Higher magnification image of the white dashed rectangle in B. I-VI, cortical layers. Brain sections were stained with CamKII antibody (for glutamatergic neurons), anti-GFP antibody (for GABAergic neurons, from a GAD67-GFP mouse line), and DAPI for nuclei visualization. (D) Higher magnification image of the yellow dashed rectangle in C. Signal for each channel was adjusted for optimal visualization. (D1) CamKII-positive neurons and GFP-positive neurons, which were never colocalized. (D2) Some jRGECO1a and nuclear Tom-positive neurons also expressed GFP (yellow cells). (D3-D6) Single channel images for each marker. Cells expressing jRGECO1a and nuclear Tom signals in nuclei (DAPI) are marked with yellow dashed circles. These cells were not CamKII-positive neurons, though some were GFP-positive neurons. (E) Percentage of GAD-positive and CamKII-positive neurons among all jRGECO1a-positive neurons. (F) Percentage of GAD-positive and CamKII-positive neurons among nuclear Tom-positive neurons. Data are presented as mean ± SEM of the original data.

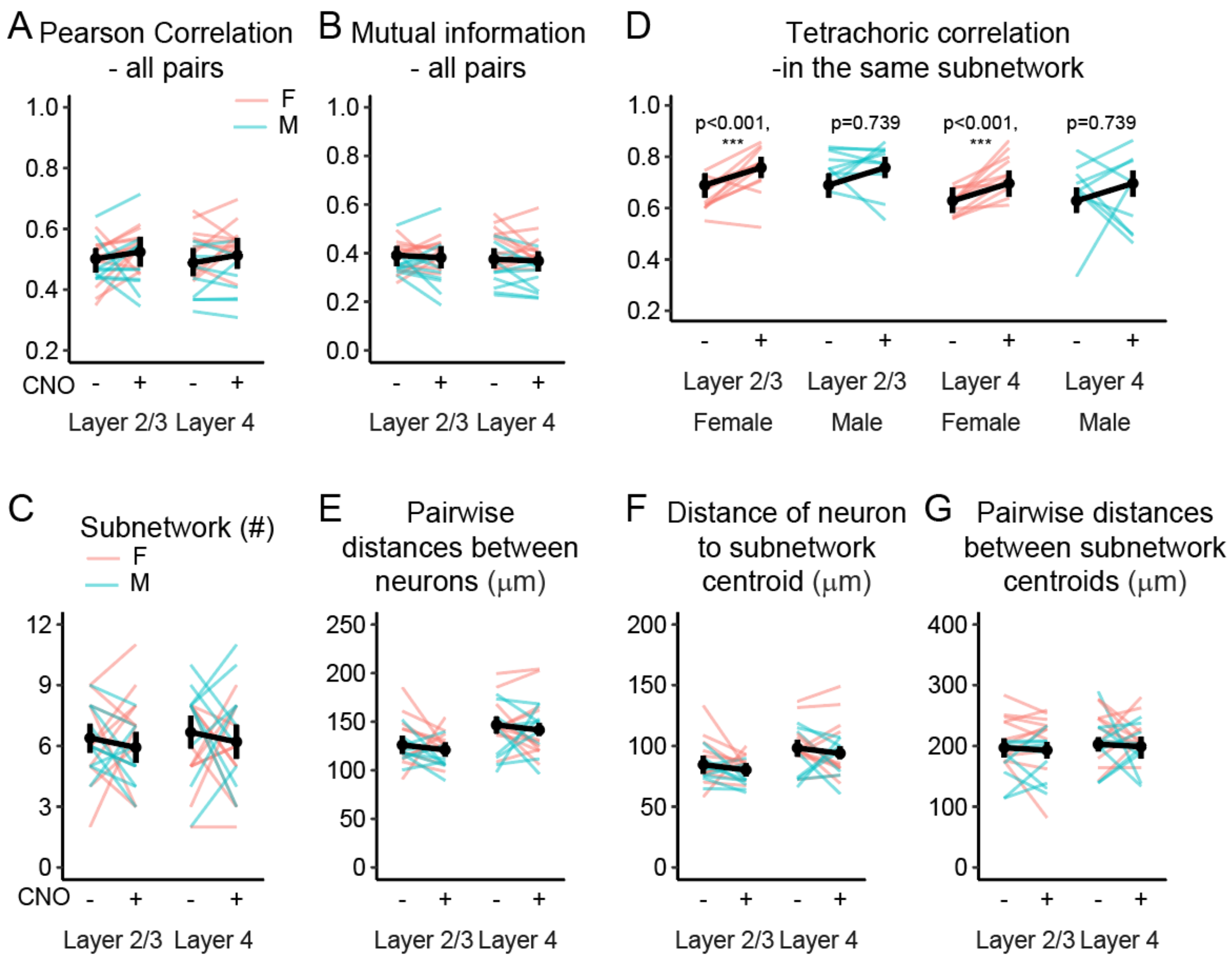


**Fig. S6**. **Enhancing GABAergic transmission did not affect correlated activity, subnetwork number, and topographic distribution of neurons within subnetworks.**

(A) Pearson correlation coefficients of neurons from all neuronal pairs. (B) Subnetwork numbers. (C) Tetrachoric correlation coefficients for neuronal pairs within the same subnetwork. (D) Pairwise distances between neurons within the same subnetwork. (E) Distances of neurons to their respective subnetwork centroids. (F) Pairwise distances between subnetwork centroids. Data are presented as mean ± 95% confidence intervals derived from the bootstrapping output.

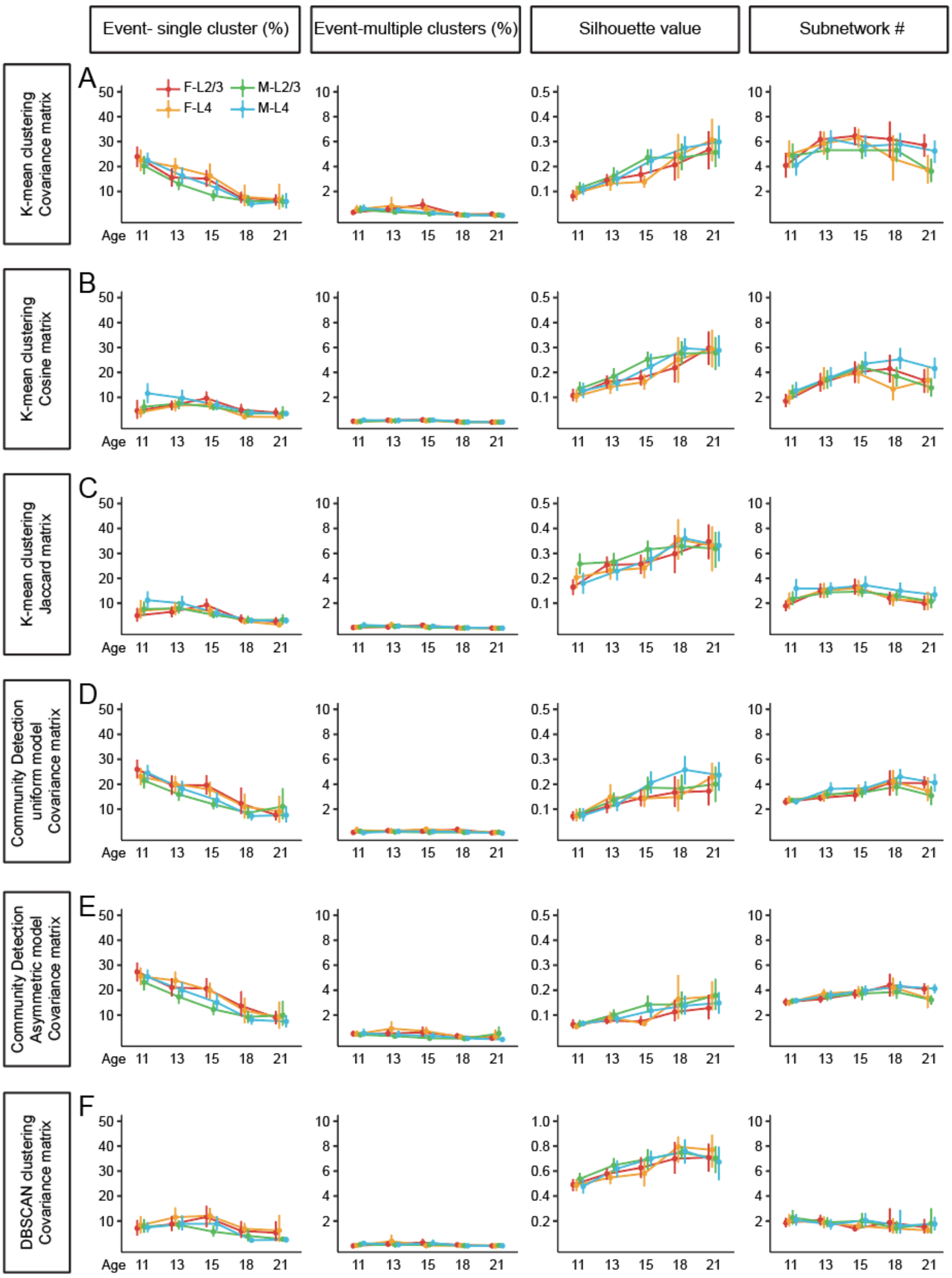

Event- single cluster (%)
Event-multiple clusters (%)
Silhouette value
Subnetwork #
K-mean clustering Covariance matrix
K-mean clustering Cosine matrix
K-mean clustering Jaccard matrix
Community Detection uniform model Covariance matrix
Community Detection Asymmetric model Covariance matrix
DBSCAN clustering Covariance matrix
A
B
C
D
E
F
F-L2/3
M-L2/3
F-L4
M-L4
Age
11 13 15 18 21

**Fig. S7. Developmental increase in silhouette value of event clusters across different clustering methods.**

(A) K-means clustering applied to the covariance similarity matrix. (B) K-means clustering applied to the cosine similarity matrix. (C) K-means clustering applied to the Jaccard similarity matrix. (D) Louvain community detection using a uniform model applied to the covariance similarity matrix. (E) Louvain community detection using an asymmetric model applied to the covariance similarity matrix. (F) DBSCAN applied to the covariance similarity matrix. Across all clustering methods, the covariance matrix consistently yielded the most optimal clustering results, identifying approximately 30% of events at P11 when using k-means clustering and community detection. All methods showed an age-dependent increase in silhouette value in both L2/3 and L4, independent of sex. Notably, k-means clustering with the covariance matrix identified a higher number of subnetworks (approximately 4) compared to other methods. Data are presented as mean ± 95% confidence intervals of the original data.

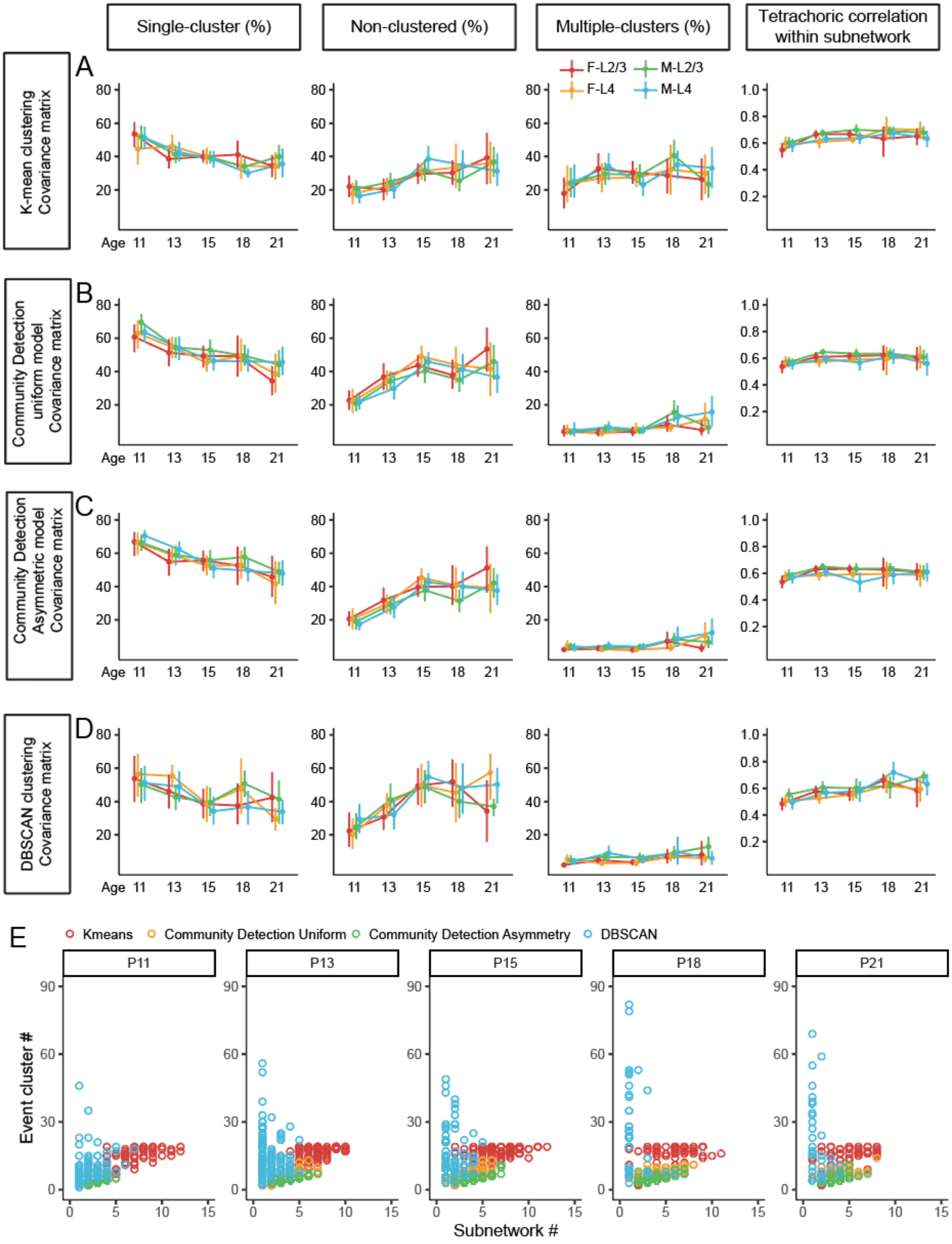
Single-cluster (%)
Non-clustered (%)
Multiple-clusters (%)
Tetrachoric correlation within subnetwork
A
K-mean clustering Covariance matrix
B
Community Detection uniform model Covariance matrix
C
Community Detection Asymmetric model Covariance matrix
D
DBSCAN clustering Covariance matrix
F-L2/3
M-L2/3
F-L4
M-L4
Age
E
Kmeans
Community Detection Uniform
Community Detection Asymmetry
DBSCAN
P11
P13
P15
P18
P21
Event cluster #
Subnetwork #

**Fig. S8. Developmental trajectory of subnetwork configuration across different clustering methods.**

(A) K-means clustering applied to the covariance similarity matrix. (B) Louvain community detection with a uniform model applied to the covariance similarity matrix. (C) Louvain community detection with an asymmetric model applied to the covariance similarity matrix. (D) DBSCAN clustering applied to the covariance similarity matrix. (E) Relationship between event cluster numbers and subnetwork numbers. Across all clustering methods, a consistent developmental trajectory in subnetwork configuration was observed. Data are presented as mean ± 95% confidence intervals of the original data.

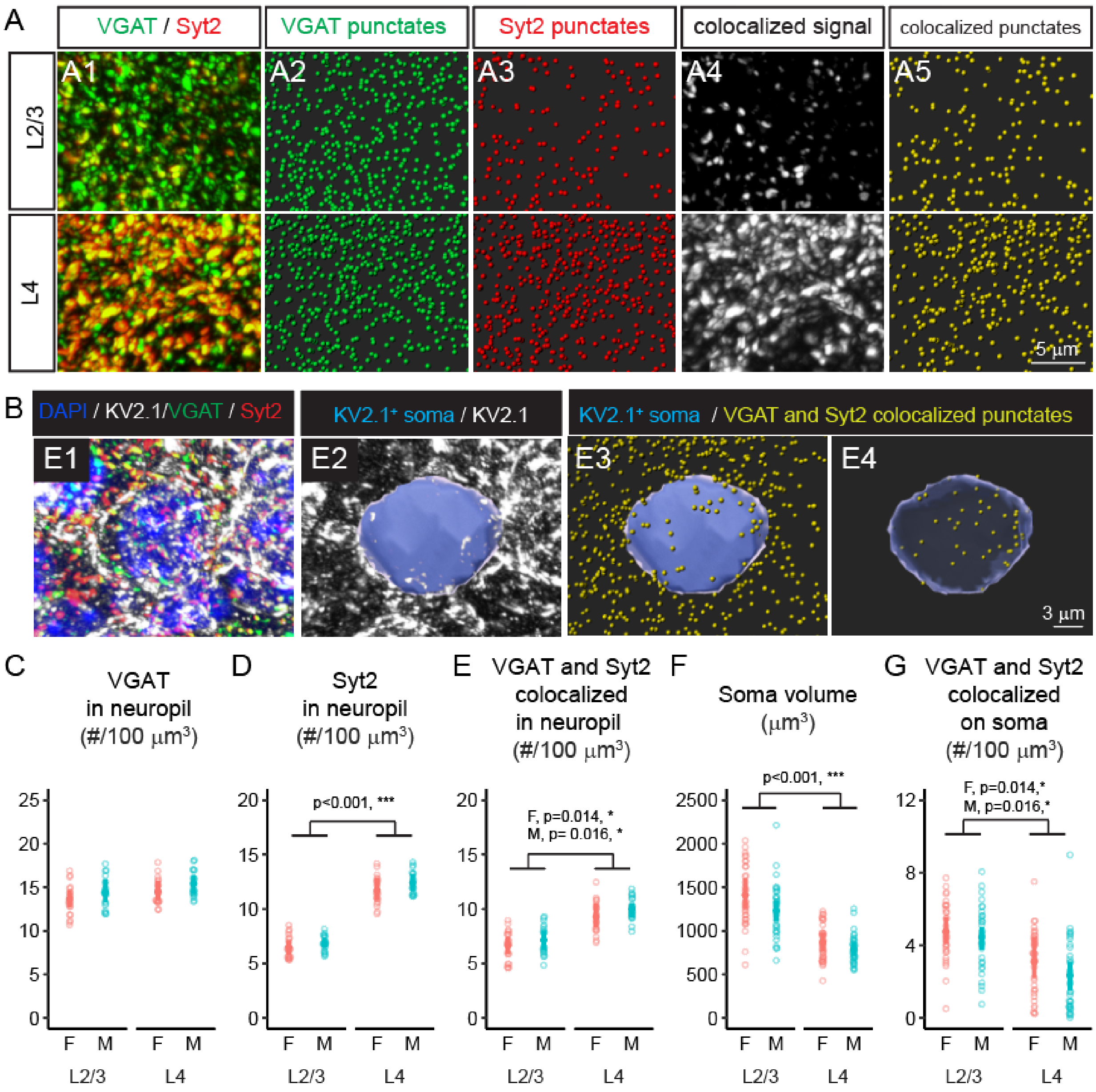


**Fig. S9. Inhibitory synapses in L2/3 and L4 of S1 cortex at P15.**

To investigate whether these sex-dependent differences might be linked to variations in inhibitory synapses, we assessed overall GABAergic input and perisomatic GABAergic input with double-staining of synaptotagmin 2 (Syt2), a marker for putative parvalbumin synapses (44) and vesicular GABA transporter (VGAT), a marker for presynaptic terminal of GABAergic synapses. (A) Confocal images showing VGAT and Syt2 signals in L2/3 and L4 (A1). Imaris spot detection was utilized to identify VGAT-positive (A2) and Syt2-positive (A3) puncta.

Colocalized VGAT and Syt2 signals were isolated using the colocalization function in Imaris (A4), and VGAT and Syt2 double-positive puncta were detected (A5). (B) Confocal images showing VGAT, Syt2, and voltage-gated K+ channel (KV2.1, visualizing soma) triple-staining to reveal potential perisomatic parvalbumin inhibitory synapses on postsynaptic soma (E1). The Imaris surface function was employed to render the somatic surface object using Kv2.1 as a reference (E2). VGAT and Syt2 double-positive puncta (as detected in A) and Kv2.1-positive somata were detected (E3). Perisomatic VGAT and Syt2 double-positive puncta within 0.25 μm of the soma were identified (E4). (C-G) Quantitative analysis showing the density of VGAT-positive puncta in the neuropil (C), Syt2-positive puncta in the neuropil (D), VGAT and Syt2 double-positive puncta in the neuropil (E), soma volume (F), and the density of VGAT and Syt2 double-positive puncta attached to the soma (G). Animal numbers: 5F (L2/3, 33 neurons; L4, 36 neurons), 5M (L2/3, 32 neurons; L4, 35 neurons). We found no significant sex-dependent differences in the density of vesicular GABA transporter $(VGAT)^+$, synaptotagmin 2 $(Syt2)^+$, or $VGAT^+/Syt2^+$ colocalized puncta in the neuropil. We noted that L4 exhibited a higher density of $Syt2^+$ and $VGAT^+/Syt2^+$ colocalized synaptic puncta compared to L2/3. Additionally, we measured perisomatic parvalbumin inhibitory synapses and found that L4 spiny stellate neurons had significantly smaller somatic volumes (F) and lower densities of these synapses compared to L2/3 pyramidal neurons (G). Data are presented as mean ± 95% confidence intervals derived from the bootstrapping output.
